# Separation of hemodynamic signals from GCaMP fluorescence measured with widefield imaging

**DOI:** 10.1101/634923

**Authors:** M.T. Valley, M.G. Moore, J Zhuang, N Mesa, D Castelli, D Sullivan, M Reimers, J Waters

## Abstract

Widefield calcium imaging is often used to measure brain dynamics in behaving mice. With a large field of view and a high sampling rate, widefield imaging can monitor activity from several distant cortical areas simultaneously, revealing cortical interactions. Interpretation of widefield images is complicated, however, by the absorption of light by hemoglobin, which can substantially affect the measured fluorescence. One approach to separating hemodynamics and calcium signals is to use multi-wavelength backscatter recordings to measure light absorption by hemoglobin. Following this approach, we develop a spatially-detailed regression-based method to estimate hemodynamics. The spatially-detailed model is based on a linear form of the Beer-Lambert relationship, but is fit at every pixel in the image and does not rely on the estimation of physical parameters. In awake mice of three transgenic lines, the Spatial Model offers improved separation of hemodynamics and changes in GCaMP fluorescence. The improvement is pronounced near blood vessels and, in contrast with other models based on regression or the Beer-Lambert law, can remove vascular artifacts along the sagittal midline. Compared to other separation approaches, the spatially-detailed model permits more accurate fluorescence-based determination of neuronal activity across the cortex.

**NEW & NOTEWORTHY:** This manuscript addresses a well-known and strong source of contamination in widefield calcium imaging data: hemodynamics. To guide researchers towards the best method to separate calcium signals from hemodynamics, we compare the performance of several commonly used methods in three commonly-used Cre-driver lines, and we present a novel regression model that out-performs the other techniques we consider.

## INTRODUCTION

Over the last few years there has been a sharp increase in the number of mouse lines with GCaMP expression throughout much of neocortex (Hasan *et al.*, 2004; Chen *et al.*, 2012; Zariwala *et al.*, 2012; Madisen *et al.* 2015; Wekselblatt *et al.*, 2016; Bethge *et al.*, 2017; Daigle *et al.*, 2018), offering the opportunity to image cortical activity with high signal-to-noise and genetically-targeted expression. As a result, there has been a resurgence in the use of widefield fluorescence calcium imaging to monitor brain dynamics *in vivo*, especially in the context of mouse behavior (Wekselblatt *et al.*, 2016; Makino *et al.*, 2017; Allen et al., 2017, Mitra et al., 2018). Widefield imaging offers a large field of view (>10 mm), enabling simultaneous imaging of almost all of neocortex, can be performed through intact skull (White et al., 2011, Silasi *et al.*, 2016), eliminating invasive craniotomies, and, compared to laser scanning techniques, widefield imaging can achieve a faster sampling rate and is relatively simple and inexpensive to implement.

Widefield imaging also presents some challenges. It lacks the optical sectioning of confocal and multiphoton microscopes, with the result that fluorescence is typically an average across cells and cellular compartments. Emission can originate from intrinsic fluorophores such as flavoproteins (Zipfel *et al.*, 2003) and fluorescence excitation and emission can be affected by endogenous absorbers, like hemoglobin. The effects of intrinsic fluorophores can be negligible in mice with bright and strongly-expressed fluorophores (e.g. Zhuang *et al.*, 2017). In contrast, the absorption of fluorescence by hemoglobin cannot be overcome with strong fluorophore expression due to the multiplicative effect of absorption on fluorescence. Furthermore, hemoglobin is a strong, broad-spectrum absorber at around 500-600 nm, which includes excitation and emission wavelengths of GCaMP. Hence changes in hemoglobin absorption greatly complicate interpretation of GCaMP fluorescence measurements, even in mice with strong GCaMP expression.

Hemodynamics encompasses multiple processes, including neuro-vascular coupling, in which neural and glial activity are accompanied by dilation of blood vessels and changes in blood oxygenation, resulting in changes in the total concentration of hemoglobin and the ratio of oxygenated to deoxygenated hemoglobin (Malonek and Grinvald, 1996; Berwick *et al.*, 2005; Hillman *et al.*, 2007; Stefanovic et al., 2008; Sirontin *et al.*, 2009; Bouchard *et al.*, 2009; Hillman et al., 2014; O’Herron *et al.*, 2016). Contraction of cardiac, pulmonary and postural muscles can also drive changes in total hemoglobin concentration, via changes in intracranial blood pressure (Gisolf *et al.*, 2004; Huo *et al.*, 2015). Postural and locomotor influences on hemodynamics are common in awake, behaving animals, making the separation of hemodynamics from activity-related changes in cellular calcium concentration a common challenge in studies of sensory-motor processing in behaving mice.

There are several strategies to separate changes in calcium indicator fluorescence from hemodynamic changes in light absorption (Bouchard *et al.*, 2009; Wekselblatt *et al.*, 2016; Allen *et al.*, 2017). One approach relies on the measurement of hemoglobin absorption using one or more diffuse reflectance or ‘backscatter’ measurements, in which some portion of the photons illuminating the brain surface undergo multiple scattering events within the tissue and return to the surface, escaping the brain to a detector. Backscatter can be measured continuously and need not conflict with fluorescence measurements, enabling continuous monitoring of hemodynamics during fluorescence imaging. From backscatter measurements, hemoglobin concentrations are often calculated using a modified Beer-Lambert relationship that relates the absorption of light by oxygenated and deoxygenated hemoglobin to the length of the mean scattering path through tissue (Bouchard *et al.*, 2009; White *et al.*, 2011; Ma *et al.*, 2016a). Mean scattering path lengths are difficult to measure empirically, forcing the use of calculated mean path lengths that do not account for local differences in scattering and absorption. Local differences in path length can lead to substantial errors in the estimated effects of hemodynamics on fluorescence measured with indicators such as GCaMPs.

Here we describe and test a spatially-detailed regression model that allows for differences in optical properties across the brain. With this model, we quantify the correction in GFP reporter mice and apply the regression model to GCaMP mice, finding that the spatially-detailed model provides improved separation of hemodynamics from changes in GCaMP fluorescence, particularly in brain areas where the separation is challenging with previous models.

## RESULTS

### Hemodynamics affect fluorescence in GCaMP-and GFP-expressing mice

We used widefield imaging to monitor fluorescence across neocortex in awake mice expressing GCaMP in neocortical pyramidal neurons. Vasculature was prominent in fluorescence images and changes in fluorescence near blood vessels were commonplace (figure 1). We observed three types of putative hemodynamic effects, defined by their spatial and temporal characteristics. First, we observed stimulus-linked changes in fluorescence that were localized within cortex (figure 1A). Fluorescence in visual cortex increased rapidly after the onset of visual stimulation followed by a prolonged sag, often to <50% peak amplitude. Following stimulus offset, fluorescence typically decreased below the pre-stimulus baseline and recovered after ∼2 seconds. The fluorescence increase is a GCaMP-mediated signal. The time course of the sag and overshoot are consistent with the ∼1 second delayed onset and 4-5 second decay of local vessel dilation during neurovascular coupling following neural activity (Malonek and Grinvald, 1996; Berwick *et al.*, 2005; Sirotin *et al.*, 2009; Ma *et al.*, 2016b). Second, we observed large amplitude (>10%) fluctuations that were restricted mainly to the midline vasculature (figure 1B), and typically included fast transient spikes in fluorescence, sometimes in bouts lasting 1-10 seconds. These fluctuations may result from changes in venous blood volume along the superior sagittal sinus that relates to movements or postural changes (Huo *et al.*, 2015, Gilad *et al.*, 2018). Finally, we observed low amplitude (<3%), global oscillations in the 8-12 Hz frequency range of the heart rate in most recordings (figure 1C). As expected, our results suggest that changes in fluorescence measured from GCaMP-expressing mice are a mixture of changes in GCaMP fluorescence and hemodynamics. In many instances, the hemodynamic effects were in the same amplitude range as changes in GCaMP fluorescence.

**Fig 1.**
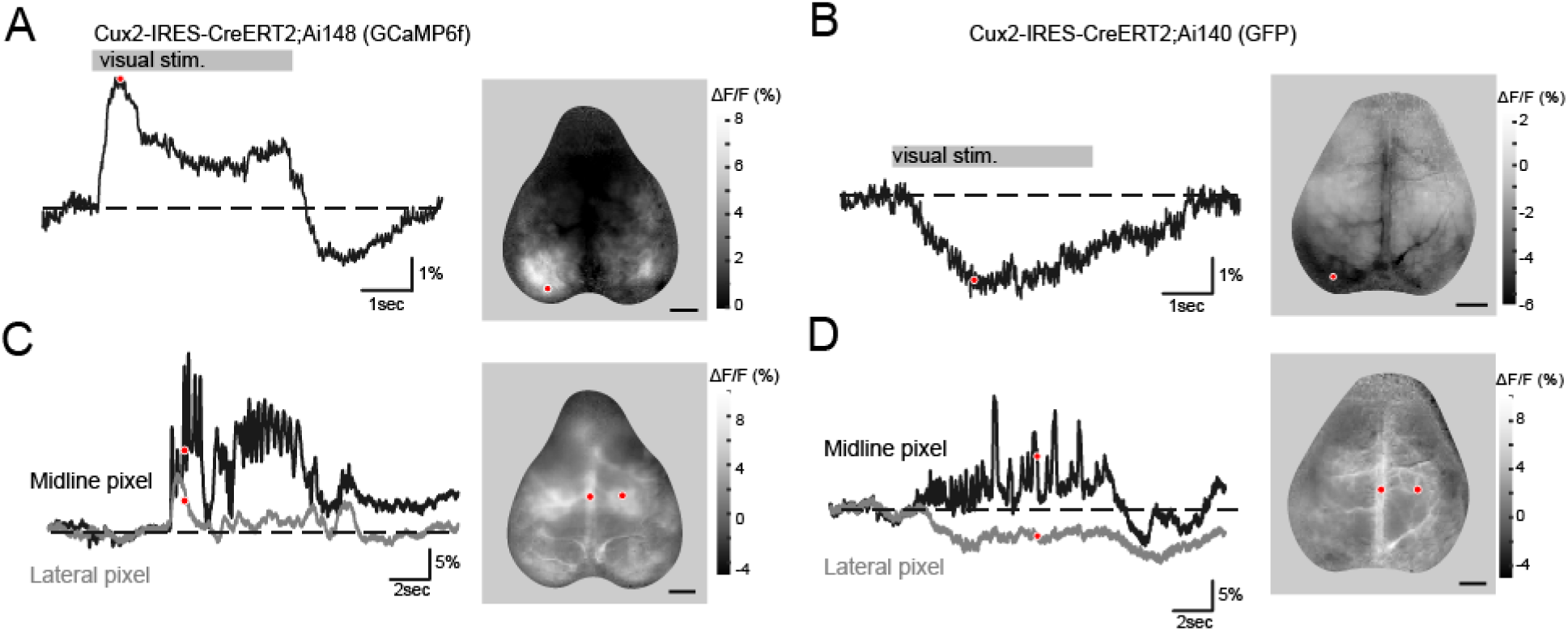
Hemodynamics in GCaMP and GFP mice. (A) Fluorescence during a five second presentation of a drifting grating. Left, fluorescence through time from a pixel in visual cortex (red marker in image). Right, image of peak fluorescence at each location. (B) Left, time-series from a Cux2-Ai140 (GFP) mouse showing mixed GCaMP and hemodynamic responses following the same stimulus. Right, fluorescence in a GFP animal 150-200 ms after stimulation. (C) Left, time-course of spontaneous activity in a GCaMP6f mouse highlighting two pixels on and off the midline vasculature. Right, image of the signal from the time indicated (red dots, left panel). (D) Same as panel C, for a GFP mouse. Image scale-bars = 2 mm.

Consistent with the suggestion that fluorescence in GCaMP animals includes a substantial hemodynamic component, we observed events with similar spatial and temporal characteristics in GFP mice (figure 1D-F), in which fluorescence does not change with intracellular calcium concentration.

### Optical strategy for simultaneous measurement of fluorescence and hemodynamics

Hemodynamics alter widefield fluorescence signals by absorbing light during excitation or emission. Absorption can be measured using backscatter (figure 2A). Hemoglobin absorption extends over a broad spectrum that includes wavelengths beyond the excitation and emission bands of GFP and GCaMP, enabling the measurement of hemoglobin concentrations using wavelengths different from those used for fluorescence (figure 2B). Furthermore, oxy-and deoxyhemoglobin absorption spectra differ substantially across the visible spectrum (figure 2B), enabling one to distinguish changes in the concentrations of oxy-(HbO) and of deoxyhemoglobin (HbR) using backscatter measurements at two wavelengths with different HbO and HbR absorption, such as 577 nm and 630 nm (Frostig *et al.*, 1990). Using two cameras, we simultaneously acquired fluorescence data at 100Hz and two backscatter wavelengths (577 nm and 630 nm) at ∼17 Hz (figure 2C). Due to these frame rates we were unable to adequately sample heart-rate hemodynamic signals (∼8-12 Hz) and used a low-pass filter to remove fluctuations at >5Hz.

**Fig 2.**
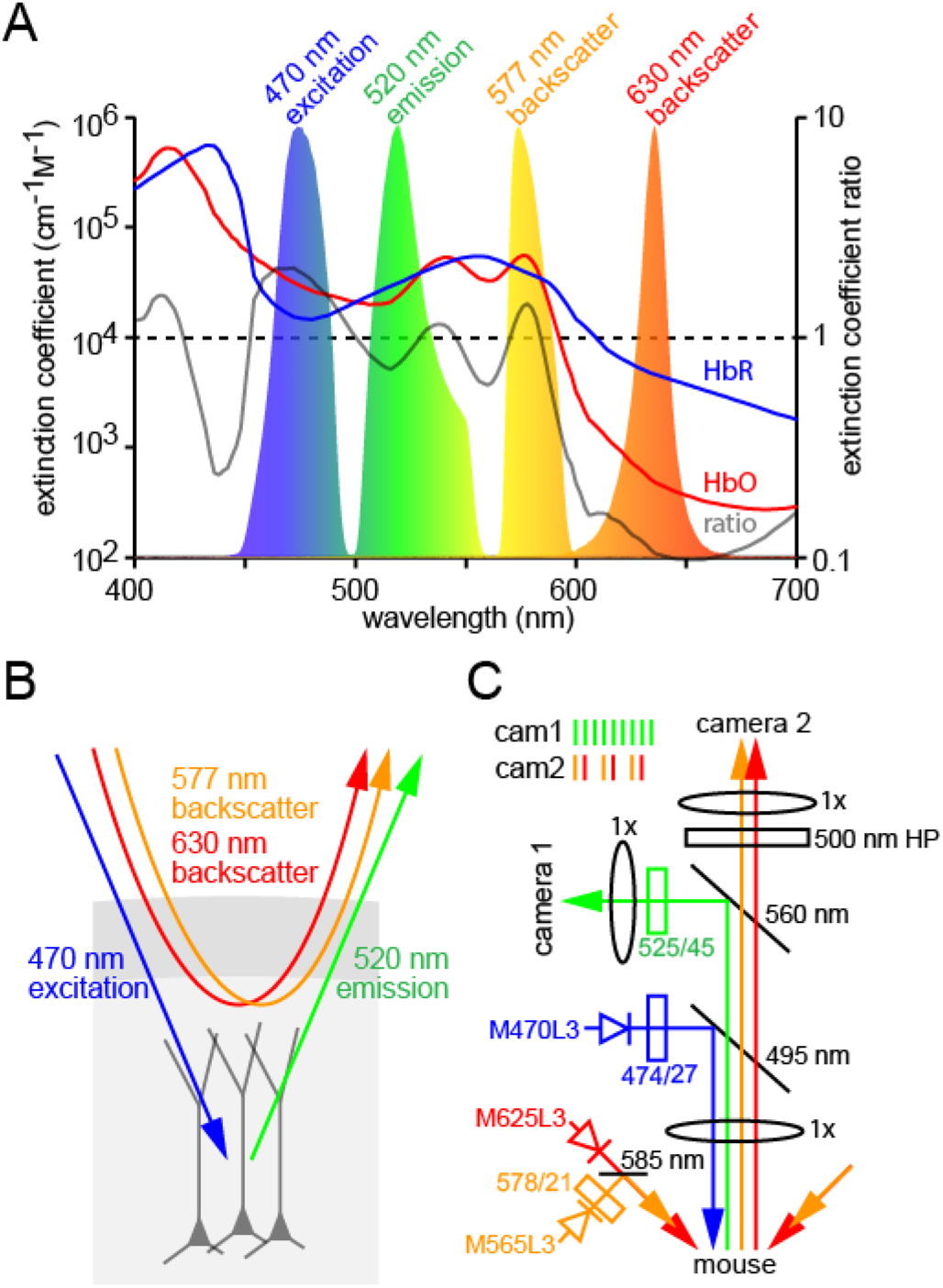
Simultaneous measurement of GCaMP fluorescence and hemodynamics. (A) Absorption spectra of oxy-and deoxyhemoglobin (HbO and HbR; left axis) in the visible spectrum, the HbO/HbR extinction coefficient ratio (right axis), and normalized spectra of wavebands used in our experiments. (B) Schematic illustration of strategy to estimate absorption of GCaMP excitation and emission (centered at ∼470 and ∼520 nm, respectively) and of illumination at ∼577 and ∼630 nm by hemoglobin. Blue illumination is susceptible to absorption when in transit from the LED to the neurons expressing GCaMP. Green fluorescence emission is susceptible to absorption while in transit from GCaMP molecules to the fluorescence camera. 577 and 630 nm illumination are susceptible to absorption during entry and during exit from the brain, before and after the scattering events that result in diffuse reflection. (C) Schematic illustration of simultaneous fluorescence and backscatter measurements, with relative locations and wavelength characteristics of LED sources, filters, lenses, dichroic mirrors and cameras (see figure S5 for a rendering of the microscope assembly). Camera frame interleaving sequence (inset, upper left) shows the continuous acquisition of fluorescence on camera 1, and strobing between detection of 577nm or 630nm backscatter and a blank frame on camera 2.

### Beer-Lambert model

The Beer-Lambert law has often been employed to estimate light absorption by oxy-and deoxyhemoglobin and thereby separate hemodynamics from changes in indicator fluorescence (Bouchard *et al.*, 2009; White *et al.*, 2011; Ma *et al.*, 2016a). The Beer-Lambert law relates absorption of light to the concentration of the absorbing species and can be used to calculate hemodynamic effects across the brain surface.

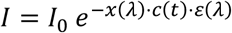

where, *I* is the measured light intensity returning from source *I*_0_, ε(λ) is the wavelength-dependent extinction coefficient of the absorbing species, *c*(*t*) is the time-varying concentration of the absorbing species, *x*(λ) is mean path length, the wavelength-dependent distance travelled by propagating light.

We calculated and removed the changes in fluorescence resulting from absorption by oxy-and deoxyhemoglobin, as described previously (Ma *et al.*, 2016a), and quantified the remaining variance. Quantifying performance in a GCaMP mouse is challenging since hemodynamics and changes in indicator fluorescence can each drive changes in measured fluorescence. Consequently, we quantified the performance of hemodynamic correction strategies in GFP-expressing Ai140 mice (Daigle *et al.*, 2019) crossed with 3 Cre lines, driving GFP expression enriched in different cortical layers: Cux2-Cre (layer 2/3), Rorb-Cre (layer 4), and Ntsr1-Cre (layer 6, figure 3C). In these three lines, GFP fluorescence accounts for ∼95% of the photons emitted from the preparation (Cux2-Cre 97.9 %, Rorb-Cre 93.7 %, Ntsr1-Cre 94.4 %, figure S1) thus we expect nearly all changes in fluorescence to result from hemodynamics with a negligible contribution from endogenous fluorophores. Complete separation of hemodynamics from changes in indicator fluorescence would reduce the variance (normalized to initial variance) to ∼0.05.

**Fig 3.**
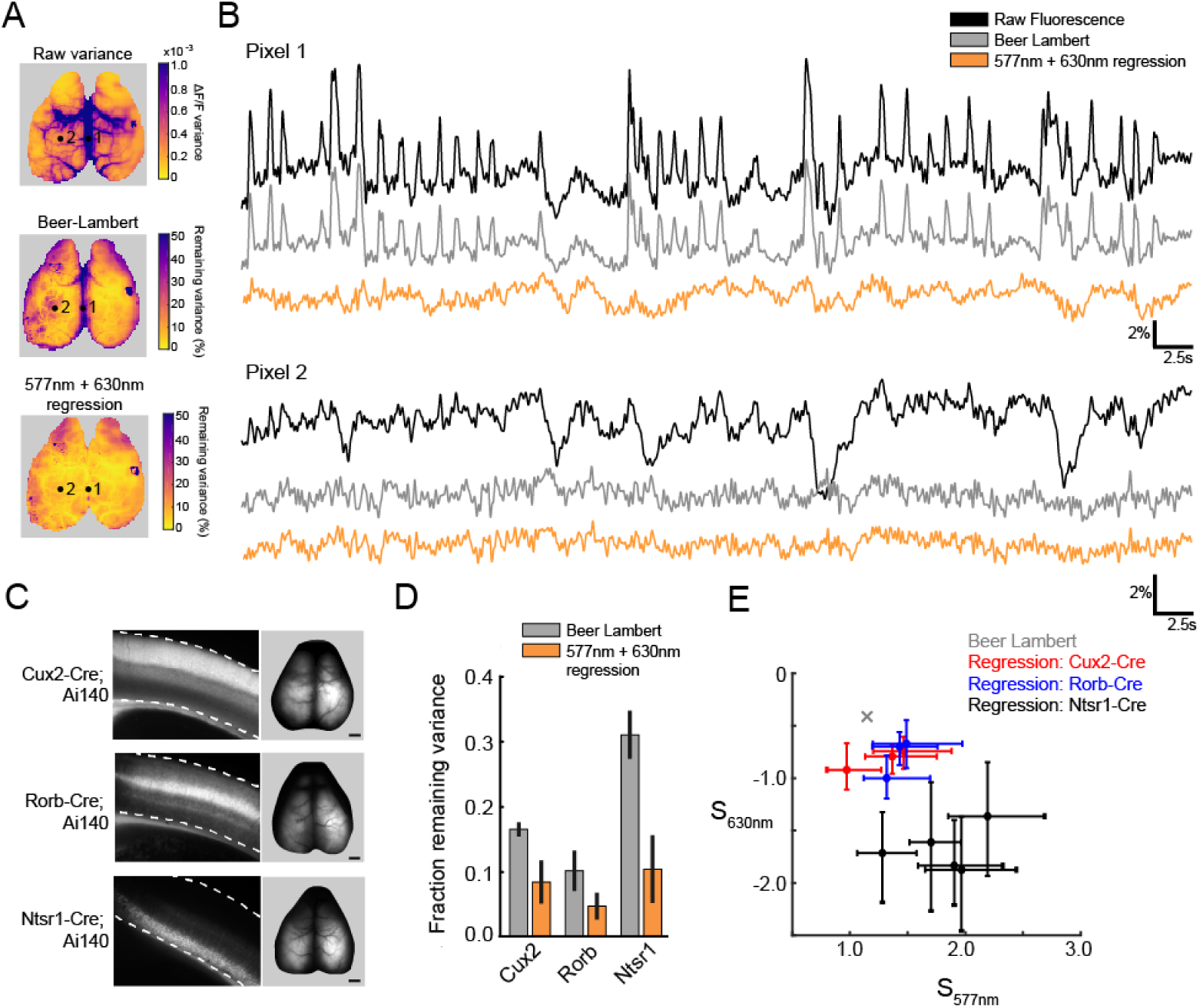
Comparison of models to explain variance in GFP mice. (A) Map of variance from a Cux2-Ai140 (GFP) mouse (top) and of remaining variance after correction using two models. Numbered dots indicate locations from which traces were extracted for panel B. (B) Example traces of fluorescence (black traces) from two example pixels (dots, panel A) and corrected fluorescence from the Beer-Lambert model (grey traces) and from dual wavelength linear regression model (yellow traces). C) Laminar expression patterns in visual cortex of Cux2-Ai140, Rorb-Ai140 and Ntsr1-Ai140 mice. Dashed lines mark the dorsal surface of cortex (top), and the ventral white-matter border of cortex. (D) Summary of variance remaining after correction (averaged across cortical surface) from three Cre lines. Bars represent mean ± STD. 3 Cux2-Ai140 mice, 3 RorbAi140 mice, 5 Ntsr1-Ai140 mice. (E) Distribution of regression coefficients for Ai140 mice, fit with 577nm and 640nm backscatter data. Using estimated parameters (path length and extinction coefficients) similar but spatially-uniform coefficients can be calculated using the Beer-Lambert relationship (grey cross, see appendix for details). Points and error bars represent median ± quartiles from all pixels in the image.

The initial variance of the fluorescence in GFP mice was not spatially uniform. Variance was ∼3-5 times greater along the midline and over large vessels than over the center of each hemisphere (figure 3A), consistent with a strong influence of movement-or posture-related hemodynamics. After Beer-Lambert correction, the variance was reduced in all locations across the brain. The median remaining variance across pixels (normalized to initial variance at each pixel), averaged across mice, was 0.17 ± 0.01 in 3 Cux2-Ai140 mice, 0.10 ± 0.03 in 3 Rorb-Ai140 mice and 0.31 ± 0.04 in 5 Ntsr1-Ai140 mice (figure 3D). However, remaining variance differed substantially with location. Normalized remaining variance was generally <0.1 in the center of each hemisphere, over anterior visual cortex and across much of somatosensory cortex (figure 3A). Performance of the correction declined towards the edges of the image, over large vessels and along the midline, where remaining variance was commonly >0.3. In these areas, it is likely that movement- or posture-related hemodynamic transients remained largely uncorrected, leaving fluorescence from midline cortical regions contaminated with hemodynamic effects.

One common simplification of the Beer-Lambert model is to choose a single extinction coefficient and path length for each wavelength band (rather than integrating over the spectrum of wavelengths; see appendix). When applied to our data, using the extinction coefficients and path lengths of the mean of each wavelength band, average remaining variance was ≥40 % greater than with the full Beer-Lambert model (remaining variance 0.24 ± 0.03 in 3 Cux2-Ai140 mice, 0.23 ± 0.04 in 3 Rorb-Ai140 mice and 0.45 ± 0.03 in 5 Ntsr1-Ai140 mice, figure S3A,D). Our results suggest that using a single extinction coefficient and path length for each wavelength band substantially impairs performance of the Beer-Lambert model and is best avoided.

### Linear regression, a spatially-detailed alternative to the Beer-Lambert model

For our Beer-Lambert calculations, we used the same path length estimates at every pixel, a necessary simplification. Naturally, performance of the model is sensitive to the choice of path lengths, particularly of the 577 nm path length (figure S2). We hypothesized that the poor performance of the Beer-Lambert model in some locations results from our inability to account for local variations in optical properties of the tissue, such as mean path length.

Two-wavelength regression is a promising alternative to the Beer-Lambert model because it may account for optical differences across the cortical surface. Unconstrained regression should always produce results that are equivalent to or better than the Beer-Lambert model because it is fit separately at each pixel, and thus has many additional degrees of freedom. Conversely, if we assume that path length is the main free parameter in the Beer-Lambert model, and if the true path lengths are identical at all locations in the tissue, then the Beer-Lambert and regression models should converge on the same solution.

We begin by establishing that, in principle, linear regression with two variables can account for hemodynamics independently for each pixel. Measured fluorescence, *I*_*F*_(*x, y, t*), can be related to two backscatter intensities, I_1_(x, y, t) and I_2_(x, y, t), using two assumptions: that the absorbance at each wavelength depends only on two hidden fluctuating variables, C_HbO_(x, y, t) and C_HbR_(x, y, t), representing the molar concentrations of oxy- and deoxy-hemoglobin, respectively; and that the absorbance on the fluorescence channel is multiplicative with the true fluorescence, *F*(x, y, t) (the fluorescence that would be evoked in the absence of absorption by hemoglobin). Hence,

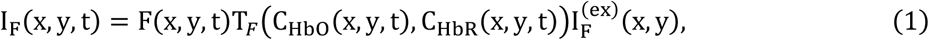

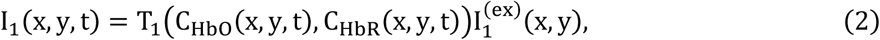

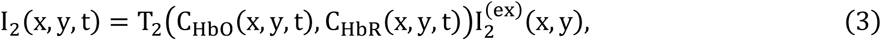

where T_*F*_, T_1_, T_2_ are transmittance functions that depend on HbO and HbR concentrations, and 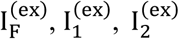 are incident intensities. The dynamic quantities relating to light intensity and hemoglobin concentration can be expressed in terms of their mean values and deviations:

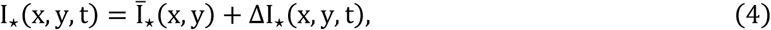

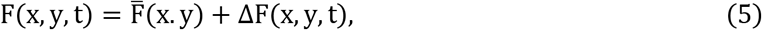

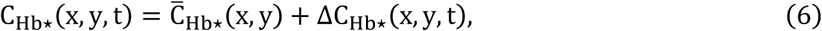

In our results, the amplitude of the fluorescence signal is large relative to its dynamic range. Under these conditions, equations 1-3 can be linearized in terms of the deviations, permitting the elimination of the hemoglobin concentrations, and enabling derivation of the regression problem (see Appendix for derivation):

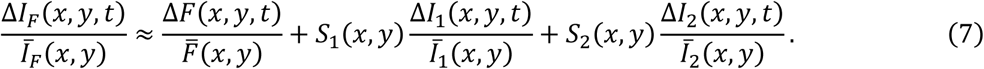

*S*_1_(*x, y*) and *S*_2_(*x, y*) are coefficient maps (figure 4C). Just as path lengths are the key unknown parameters in the Beer-Lambert model, regression coefficients are the key to separating hemodynamics from changes in indicator fluorescence using the regression model. As a result of the linearization, regression coefficients can be calculated from path lengths (figure 3E, and see Appendix).

**Fig 4.**
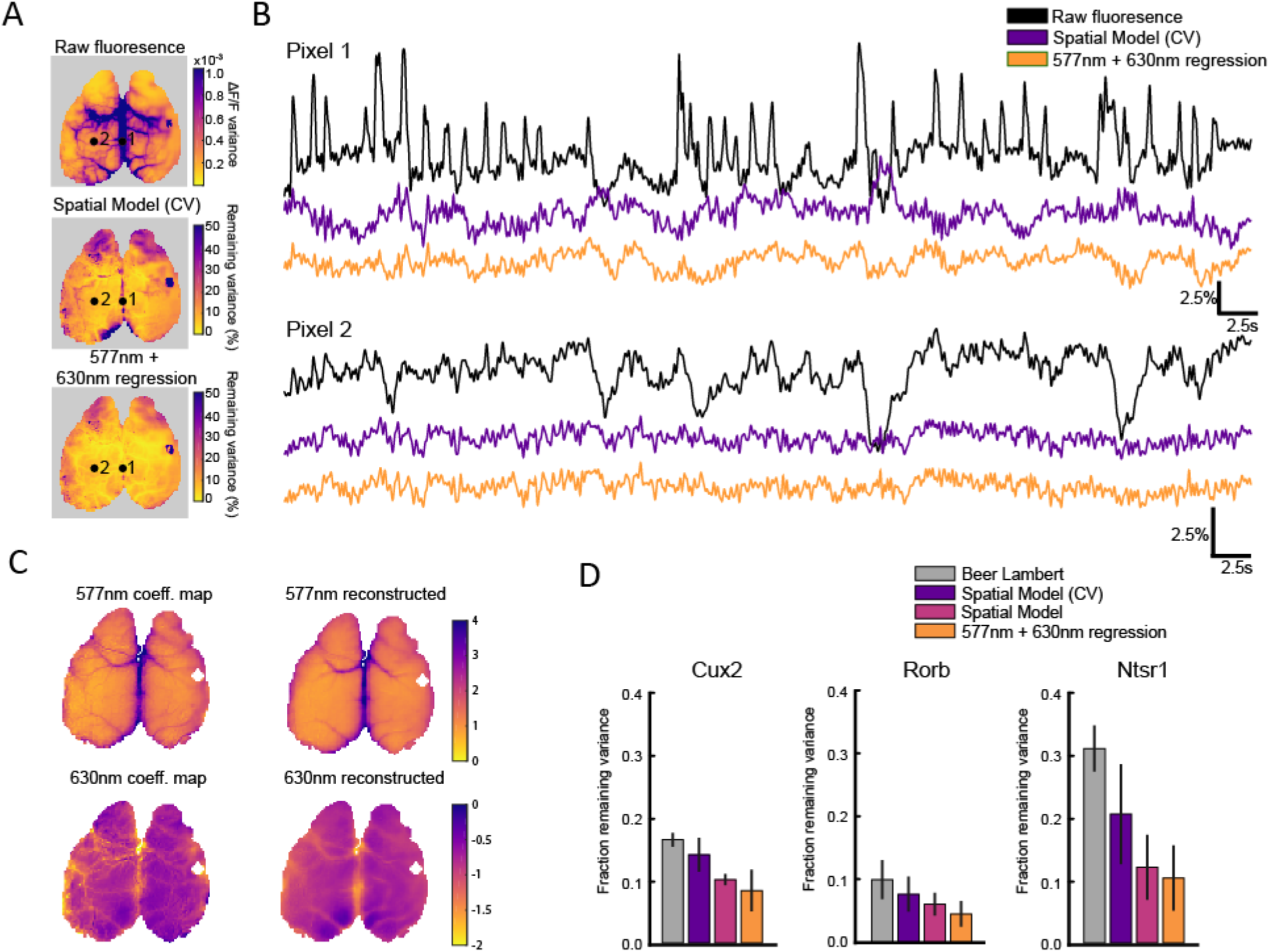
Evaluation of the Spatial Model. (A) Map of variance from a Cux2-Ai140 (GFP) mouse (top) and of remaining variance after correction using the Spatial Model using averaged parameters generated during across-animal leave-one-out cross-validation (Spatial Model CV), and the two-wavelength regression model. Numbered dots indicate locations from which traces were extracted for panel B. (B) Example traces of fluorescence (black traces) from two example pixels (dots, panel A) and of corrected fluorescence from Spatial Model CV and from dual wavelength linear regression model (yellow traces). (C) Left, coefficient maps generated using spatially-detailed regression of GFP fluorescence onto 577 nm and 630 nm backscatter. Right, reconstructions of coefficient maps (from held-out data) using the Spatial Model. (D) Comparison of remaining variance across models. Mean ± standard deviation across all animals, where the median remaining variance is taken across all pixels for each animal.

In GFP mice, where changes in measured fluorescence result only from hemodynamics, Δ*F*(t) ≈ 0 permitting simplification of equation 7:

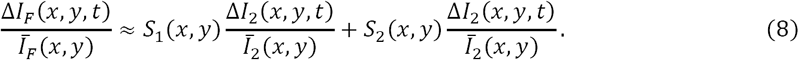

From equation 8, we can solve for the coefficients (S_1_ and S_2_) by pixel-wise linear regression of ΔI_*F*_/Ī_*F*_ onto ΔI_1_/Ī_1_ and ΔI_2_/Ī. The effectiveness of this approach is evaluated via the remaining fluorescence intensity, *F*(x, y, t)

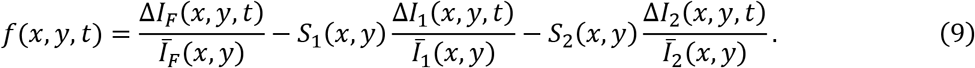

f(x, y, t) is expected to approximate 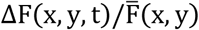, so that for GFP mice *F* ≪ ΔI_*F*_/ Ī_*F*_ ≈ 0. Hence, linear regression with two variables can account for fluorescence variance due to hemodynamics independently for each pixel, in the absence of calcium-dependent variance. Changes in the corrected fluorescence decline towards zero as performance of the regression model approaches the limit of complete separation.

When applied to GFP fluorescence, the regression model improved on the performance of the Beer-Lambert model. Averaging across all pixels, the median remaining variance across pixels, averaged across mice, was 0.08 ± 0.03 in 3 Cux2-Ai140 mice, 0.05 ± 0.02 in 3 Rorb-Ai140 mice and 0.10 ± 0.05 in 5 Ntsr1-Ai140 mice (figure 3D), ∼60-70% reduction in remaining variance relative to the Beer-Lambert model. Regression produced uniformly good correction across most of the brain, with the improved performance being particularly noticeable over large vessels, along the midline and towards the edges of the brain, where the Beer-Lambert correction was poor (figure 3A). The large, fast putative movement-or posture-related hemodynamic transients along the midline were largely eliminated by spatially-detailed regression (figure 3B). We conclude that pixelwise regression using two backscatter measurements offers improved separation of hemodynamics from changes in indicator fluorescence, likely because regression enables the model to account for differences in optical properties across the brain.

Interestingly, the mean regression coefficients required for optimal correction differed between mouse lines. Regression coefficients for Cux2-Ai140 and Rorb-Ai140 were clustered (figure 3E), perhaps because of their common GFP expression in pyramidal apical dendrites that ramify through layers 1-3. Coefficients were noticeably different for Ntsr1-Ai140 mice, with expression in layer 6 pyramidal neurons. These differences are consistent with the idea that optical properties such as fluorescence mean path lengths differ between mouse lines and need to be adjusted for optimal correction.

### Single-wavelength linear regression

Single-wavelength regression has often been used to separate hemodynamics from changes in indicator fluorescence. Generally, single-wavelength regression employs backscatter at an isosbestic wavelength such as 530 nm or 577 nm. Consequently, one might expect single-wavelength regression to best account for changes in total hemoglobin concentration (Frostig *et al.*, 1990) but not changes in blood oxygenation. Again, we used the same data set to compare the performance of single- and two-wavelength regression.

We regressed the 577 nm backscatter measurement against GFP fluorescence, pixelwise:

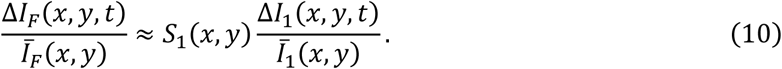

The remaining variance after 577nm single-wavelength regression was 0.13 ± 0.06 in 3 Cux2-Ai140 mice, 0.09 ± 0.03 in 3 Rorb-Ai140 mice and 0.24 ± 0.09 in 5 Ntsr1-Ai140 mice (figure S3), ∼2-3 times more remaining variance than two-wavelength regression and comparable to the Beer-Lambert model. Unlike the Beer-Lambert model, 577 nm single-wavelength regression performed reasonably well across most pixels (figure S3A), underlining the importance of allowing for differences in optical properties by tuning the model at each pixel. Overall, the performance of 577 nm single-wavelength regression was intermediate between that of the Beer-Lambert and two-wavelength regression models.

A simplified 577 nm single-wavelength regression model using a coefficient of 1 at all pixels (Xiao *et al.*, 2017) left remaining variance of 0.20 ± 0.07 in 3 Cux2-Ai140 mice, 0.21 ± 0.03 in 3 Rorb-Ai140 mice and 0.43 ± 0.03 in 5 Ntsr1-Ai140 mice (“577nm Ratiometric Demixing”, figure S3). Like other models not tuned pixelwise, correction was particularly poor over large vessels, along the midline and towards the edges of the hemispheres. However, pixelwise regression is no guarantee of performance: 630 nm single-wavelength pixelwise regression offered poor performance with remaining variance of 0.84 ± 0.10 in 3 Cux2-Ai140 mice, 0.61 ± 0.04 in 3 Rorb-Ai140 mice and 0.68 ± 0.15 in 5 Ntsr1-Ai140 mice (figure S3).

### A Spatial Model to predict regression coefficients in GCaMP mice

Tuning a regression model at each pixel improves performance in many brain areas, but how can we generate pixel-wise coefficient maps in GCaMP mice? In GFP mice, we found coefficients by regressing changes in backscatter intensity against changes in fluorescence at each pixel, under the assumption that changes in fluorescence were due to hemodynamics. This assumption is not valid for GCaMP mice because hemodynamics correlate with neuronal activity, so direct regression is not an option. Instead we developed a ‘Spatial Model’ that predicts regression coefficients at each pixel using features of backscatter and fluorescence images that can be measured in GCaMP mice. We trained and validated the performance of this Spatial Model on movies from GFP mice and applied the model to GCaMP mice.

The Spatial Model uses several statistical projections (e.g. stdev, skew or kurtosis) of backscatter and fluorescence images to predict coefficient maps (figure 4C). We regressed primary coefficient maps from GFP animals onto 19 statistical projections (figure S4) to determine the weighting of features (table S1) that best predicted the regression coefficients. We tested the performance of the Spatial Model in 2 stages. First, we trained and tested spatial weights from the same GFP mouse to reveal the performance limit of the Spatial Model. Remaining variance, averaged across all pixels, was 0.14 ± 0.03 in 3 Cux2-Ai140 mice, 0.08 ± 0.03 in 3 Rorb-Ai140 mice and 0.21 ± 0.08 in 5 Ntsr1-Ai140 mice (figure 4A), close to the performance of the regression model. Second, we used leave-one-animal-out cross-validation to generate mean spatial weights for each mouse line (table S1) and applied these mean spatial weights to generate coefficient maps and separate hemodynamics and changes in indicator fluorescence in the left out (test) mouse. This second test replicates the procedure that will be later employed in GCaMP mice in which spatial weights estimated from GFP mice are applied to GCaMP mice. The median remaining variance of all pixels in this second test, averaged across animals, was 0.14 ± 0.03 in 3 Cux2-Ai140 mice, 0.08 ± 0.03 in 3 Rorb-Ai140 mice and 0.21 ± 0.08 in 5 Ntsr1-Ai140 mice. Hence in all three mouse lines, the remaining variance was greater than with the two-wavelength regression model (that cannot be directly applied to GCaMP mice) but less than the remaining variance of the Beer-Lambert model (that can be applied to GCaMP mice). Much of the improvement in performance of the Spatial Model over the Beer-Lambert model is near vessels and along the midline. Hence, we expect the Spatial Model to offer improved separation of hemodynamics and changes in indicator fluorescence versus the Beer-Lambert model when applied to GCaMP mice, particularly for cortical areas along the midline and near large vessels.

### Separation of hemodynamics from changes in indicator fluorescence in GCaMP mice

We used the spatially-detailed regression model to separate hemodynamics from changes in indicator fluorescence in Cux2-Ai148 mice, with GCaMP6f in superficial pyramidal neurons. We trained the Spatial Model on GFP mice and then applied the resulting coefficient maps to genetically-matched GCaMP6 mice (Cux2-Ai140 vs Cux2-Ai148 mice, etc.). During spontaneous activity, variance was sometimes increased, and at other times reduced, especially during fast changes in fluorescence during events along the midline (figure 5A). During presentation of a visual stimulus, the vasodilatory sag was reduced, in GFP (figure 5B) and in GCaMP mice (figure 5C) and the baseline overshoot was eliminated. In the GCaMP results, the difference between original and corrected traces was similar in amplitude and time course to the change in measured fluorescence in GFP mice, consistent with successful separation of hemodynamics and changes in indicator fluorescence in GCaMP mice.

**Fig 5.**
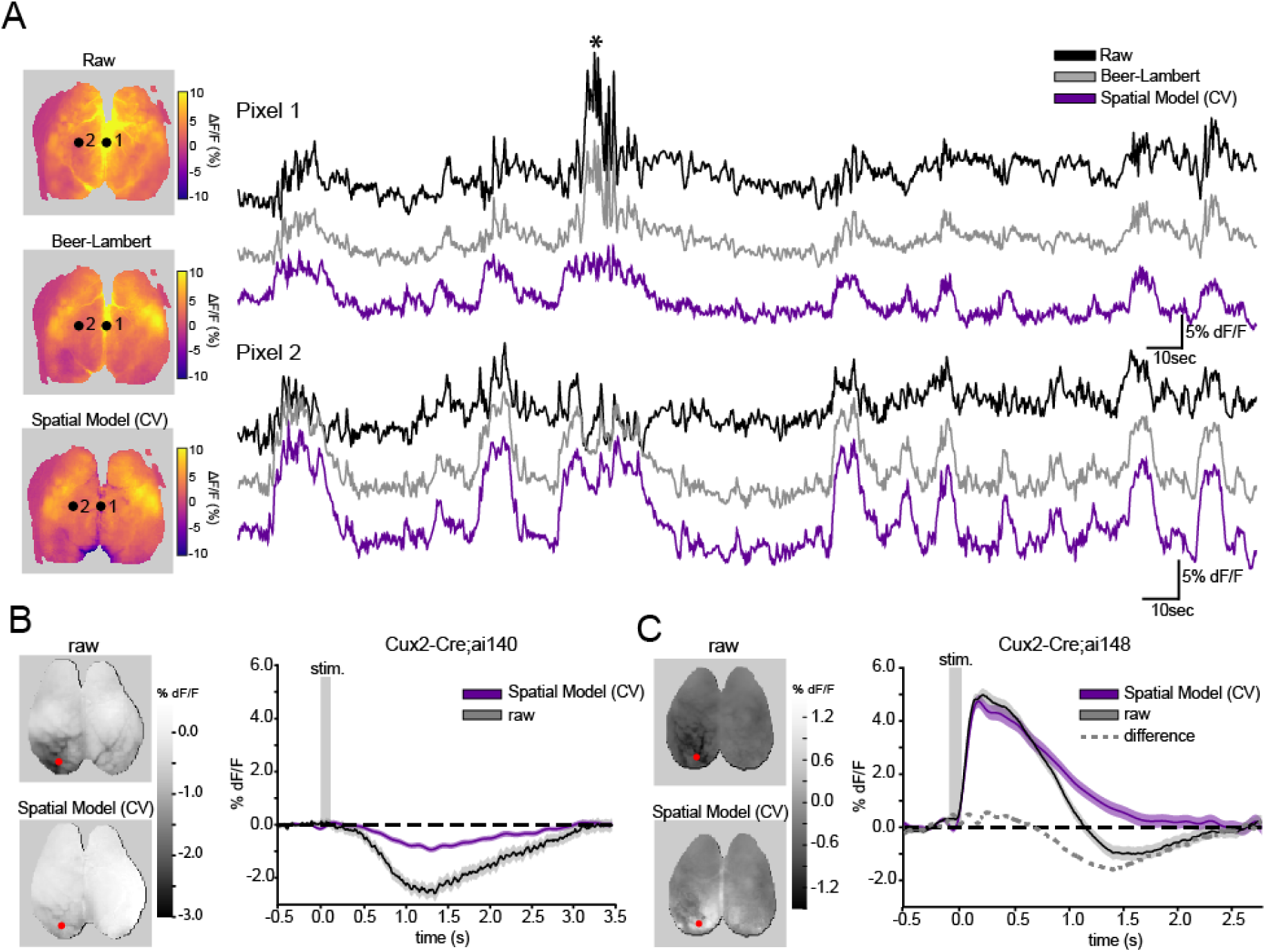
Demixing hemodynamic responses to a visual stimulus in GFP and GCaMP mice. (A) Spontaneous activity from a Cux2-Ai148 (GCaMP6f) mouse and correction with Beer-Lambert Model and Spatial Model. Left, images of fluorescence at a single time point, during midline activity (asterisk, traces). Right, ΔF/F traces for two pixels (images, left) before (black) and after correction with Beer-Lambert Model (grey) and Spatial Model (purple). (B) Left, map of mean ΔF/F 1.5-2 seconds post stimulus. 100 ms flashed visual stimulus to the right eye. Cux2-Ai140 (GFP) mouse. Right, raw trace (black) and trace after Spatial Model correction (purple) extracted from a pixel in V1 (red dot, left). (C) Left, map of mean ΔF/F 1.5-2 seconds post stimulus. 100 ms flashed visual stimulus to the right eye. Cux2-Ai148 (GCaMP6f) mouse. Right, raw trace (black) and trace after Spatial Model correction (purple) extracted from a pixel in V1 (red dot, left).

## DISCUSSION

We quantified the effects of hemodynamics on fluorescence measured from GFP mice with widefield fluorescence imaging and the performance of several models in separating hemodynamics from changes in indicator fluorescence. Previous methods have used physical models of light scattering and absorption (Malonek and Grinvald, 1996; Berwick *et al.*, 2005; Hillman *et al.*, 2007, Devor *et al.*, 2012), leveraging research on tissue absorbance spectra (Takatani and Graham, 1979; Wray *et al.*, 1988), endogenous fluorophores (Zipfel *et al.*, 2003), and models of scattering in tissue (Kohl *et al.*, 2000). Correction strategies based on physical parameters, such as the Beer-Lambert model, may overgeneralize and not account for unique features in cranial windows, or tissue heterogeneities such as surface vasculature. One might expect separation models based on physical parameterization to perform poorly in some locations. One advantage of regression models is that they can account for pixelwise differences in optical properties. We found that the Beer-Lambert model removed most, but not all hemodynamic effects and poor separation was common near surface blood vessels. Regression against two backscatter wavelengths out-performed the Beer-Lambert model, with outperformance being particularly noticeable near blood vessels. Pixelwise tuning of the model was likely the basis for this outperformance.

Central to our approach is the linearization of the Beer-Lambert equation. The resulting linear model cuts through the complexity of trying to estimate the spectral integrals of path lengths and extinction coefficients of the Beer Lambert equation by simplifying the problem to a linear estimation of two variables.

The performance of single-wavelength regression was similar in some respects to that of the Beer-Lambert model. Both performed adequate correction at many pixels, but poorly near blood vessels. The Beer-Lambert model requires two backscatter measurements and, therefore, either two backscatter cameras or multiplexed measurements on a single camera, making single-wavelength regression considerably easier to implement.

Regression against two backscatter wavelengths out-performed other models, eliminating >90% of variance in most GFP mice. In GCaMP mice, using the Spatial Model, residual variance was ∼0.1-0.2. Hence correction is not perfect, but is close to the expected limit of ∼0.05 remaining variance after removal of all hemodynamic effects. Residual variance ∼0.1 above the expected limit corresponds to ∼95% reduction in the amplitude of the mean hemodynamic transient (0.9^1/2^), meaning that a large midline hemodynamic artifact that initially caused a 10% change in apparent ΔF/F in a GCaMP mouse would be attenuated to ∼0.5% ΔF/F after application of the Spatial Model. The Spatial Model does not completely remove hemodynamic effects from fluorescence traces in GCaMP mice, but typically reduces the amplitudes of apparent changes in fluorescence to close to the noise of GCaMP measurements.

One important caveat is that we considered hemodynamic variance at <5 Hz, to avoid aliasing the heart-rate oscillations at ∼8-12 Hz. One result is that fast transients often observed along the midline were not always fully corrected by the Spatial Model. We fully expect that midline transients and heart-rate signals would be effectively corrected by our method by pushing camera acquisition to higher frame rates. The method to build statistical maps of backscatter data in the Spatial Model could also potentially be improved by using alternate methods for spatial feature extraction such as using deep-learning techniques trained on the primary coefficient maps. Similarly, it is possible that non-linearities, such as fitting time-varying regression coefficients, could form the basis for future improvements in hemodynamic demixing, but the margin for further improvement appears small since the Spatial Model accounts for most hemodynamic effects on fluorescence.

An alternative to using backscatter measurements is to stimulate GCaMP with ultra-violet light (∼410 nm) where GCaMP fluorescence is not calcium-dependent. This approach directly measures and corrects for changes in GCaMP emission (but not excitation), presumably caused by hemodynamics. We found this strategy to be effective over short time periods, but phototoxicity is a concern with ultra-violet illumination. In addition, the very shallow penetration of 410nm light (estimated mean scattering path length is ∼8µm) may limit the performance of this approach for correcting for the absorption of fluorescence from deep in tissue.

We quantified model performance in GFP mice because we lack the tools to measure the ground truth calcium signal at all cortical locations simultaneously in GCaMP mice. We then developed the Spatial Model to permit separation of hemodynamics and changes in indicator fluorescence. Our strategy used genetically-matched GFP and GCaMP mice, made no assumptions regarding the source of the fluorescence variance, and was effective when applied to several GCaMP lines. Our strategy has drawbacks. It requires experiments in GFP and GCaMP mice with matched expression and, lacking ground truth, it cannot provide a direct measure of separation performance.

We propose that experimenters consider different strategies for minimizing the effects of hemodynamics on widefield fluorescence measurements. The use of longer-wavelength indicators, such as RCaMPs, will reduce the effects of hemodynamics, but GCaMP indicators are in more widespread use. With blue-green indicators, options include the Beer-Lambert model, single-wavelength regression and our Spatial Model. Which offers the optimal balance of performance and experimental complexity will depend on several factors, including the proximity of the cortical areas of interest to blood vessels, the extent to which behavior and hemodynamic effects on fluorescence are coupled (best measured in GFP mice), the number of cameras available and extent to which they can be synchronized, the frequency band in which the fluorescence signals of interest occur, and the availability of transgenic or viral resources to produce matched GFP and GCaMP expression.

## METHODS

### Animals and surgical preparation

We used six mouse lines, from crossing three Cre driver lines with two reporter lines.

Cre lines:

Rorb-IRES-Cre, RRID:IMSR_JAX:023526, Harris *et al.* (2014).
Cux2-CreERT2, RRID:MMRRC_031778-MU, Franco *et al.* (2012).
Ntsr1-Cre_GN220, RRID:MMRRC_030648-UCD, Gong *et al.*, (2007).

Reporter lines:

Ai140, RRID:IMSR_JAX:030220, Daigle *et al.* (2018).
Ai148, RRID:IMSR_JAX:030328, Daigle *et al.* (2018).

Crosses were made between animals hemizygous for ai140 or ai148, and animals that were either hemi or homozygous for Cre. We refer to these crosses using abbreviations: Cux2-Ai140, Rorb-Ai140, Ntsr1-Ai140, Cux2-Ai148, Rorb-Ai148, Ntsr1-Ai148.

For Cux2-CreERT2 animals, tamoxifen was administered via oral gavage (50 mg/ml in corn oil) at 0.2 mg/g body weight for 3-5 days. Mice were used for experiments a minimum of two weeks following induction.

Wide-field imaging was performed on 7-30 week-old male and female mice through the intact skull using a modification of the method of Silasi *et al.* (2016). Under isoflurane anesthesia, the skull was exposed and cleared of periosteum, and a #1.5 borosilicate coverslip (Electron Microscopy Sciences, #72204-01) was fixed to the skull surface with a layer of clear Metabond (Sun Medical Co.). A 3D-printed light shield was fixed around the coverslip using additional Metabond, and the outward-facing surfaces were coated with an opaque resin (Lang Dental Jetliquid, MediMark). A custom titanium headpost was fixed posterior to the lightshield/coverslip and dorsal to the cerebellum using Metabond.

Animal experiments were performed in accordance with the recommendations in the Guide for the Care and Use of Laboratory animals of the National Institutes of Health. All animals were handled according to institutional animal care and use committee protocols of the Allen Institute for Brain Science, protocol numbers 1408 and 1705.

### Image acquisition and initial image processing

Mice were head-restrained and free to run on a 16.5 cm diameter disk. With exception of experiments with visual stimulation, mice were in the dark and spontaneously active. Visual stimuli were displayed on a 27” LCD monitor placed 13.5 cm from the right eye and consisted of high-contrast gabor gratings spanning 20 degrees of the visual field, positioned at the center of the visual field. Images were presented at 0.25 Hz and averages were made from a minimum of 50 stimulus presentations.

Images were produced by a tandem-lens macroscope of custom optomechanical design (figure S5) built around a pair of identical lenses (Leica 10450028). Epifluorescence illumination used a 470 nm LED (Thorlabs M470L3) filtered (Semrock FF01-474/27-50) and reflected by a dichroic mirror (Semrock FF495-Di03-50 × 70) through the objective lens. Backscatter illumination in yellow used a LED (Thorlabs M565L3) and a bandpass filter (Semrock F01-578/21), and backscatter illumination in red used a LED (Thorlabs 625L3). Yellow and red illumination was focused onto a 1-to-7 fan-out fiber bundle (Thorlabs BF72HS01), and the termination of each of the seven fibers was uniformly spaced circumferentially around a custom light shield surrounding the imaging objective with each fiber terminating at 45 degrees incident to the brain surface. Fluorescence emission was separated from the two backscatter wavelengths using a dichroic beamsplitter (Semrock, FF560-FDi01-50×70) and passed through an emission filter (Semrock FF01-525/45-50) to a camera while backscatter passed through a high-pass filter (Edmund Optics Y-50, 500nm) to a second camera.

Image acquisition used two Hamamatsu Flash4.0 v3 sCMOS cameras. One camera used for detection of fluorescence operated with 10 ms rolling shutter exposure (100 Hz), and the second which detected backscatter received triggered exposures at 50Hz. We used 4 illumination and detection wavelength bands: fluorescence excitation, fluorescence emission, backscatter at ∼577 nm and backscatter at ∼630 nm. We measured the spectrum of each band with a spectroradiometer (SpectroCAL, Cambridge Research Systems). The mean wavelengths of our four bands were 472, 522, 577 and 630 nm. Backscatter measurements centered at 577 and at 630 nm were interleaved with a blank frame (for calculation of fluorescence bleed-through) and thus the final sample rate for each channel on the backscatter camera was 50/3 ∼= 17 Hz.

Analysis was performed using Matlab (v2018a, Mathworks) or Python 2.7. Images were spatially down-sampled from 2048 × 2048 to 128 × 128 pixels by averaging. A camera offset of 100 counts was subtracted and camera counts converted to photo-electrons (2.19 counts per photo-electron). Backscatter signals, acquired at 17 Hz, were filtered with a 5 Hz Butterworth low pass filter to prevent aliasing of the heart beat and were then up-sampled to 100Hz, and spatially and temporally aligned to the 100 Hz fluorescence signal.

Calculation of remaining variance was made with pixelwise normalization to initial variance. To account for its skewness, we took the median of the distribution of remaining variance across pixels to represent the remaining variance in each mouse and averaged across all mice of a genetic background. In GFP mice, the shot noise floor was substantially reduced by the spatial averaging and was 0.5 % of the initial variance.

### Beer-Lambert model

For Beer-Lambert calculations, we obtained extinction coefficients from tabulated hemoglobin spectra (http://omlc.ogi.edu/spectra/hemoglobin/summary.html), and path lengths from Monte Carlo simulations (Ma *et al.*, 2016a). For each wavelength band, we numerically solved the integral of the path lengths, using the measured spectra of the fluorescence and backscatter wavelength bands (Appendix, equation 11), and tabulated GFP excitation and emission spectra (http://www.tsienlab.ucsd.edu/Documents.htm). The approximations used in what we term the Beer-Lambert, and Simplified Beer-Lambert models are described in detail in the appendix.

### Regression model

We used either 577nm, 630nm or both channels of backscatter data and calculate the weights at each pixel that best fit fluorescence from a GFP-expressing animal using ordinary least-squares. Alternatively, these primary weights were estimated using the Spatial Model which consists of 19×2 secondary weights (table S1) generated by separately fitting 19 statistical projections of backscatter and fluorescence data (see figure S4) onto the 577nm and 630 primary weightings. A detailed examination of the relationship between the Beer-Lambert and regression models is given in the appendix.

Matlab code to train and test Spatial Models, and perform all variants of regression and Beer-Lambert demixing is available online: https://github.com/MichaelGMoore/MultiChanHemo

## ACKNOWLEDGEMENTS

We would like to thank Jesse Miles for helpful technical assistance in early stages of this project, and we would like to thank Kevin Takasaki, Nicholas Steinmetz, Elizabeth Hillman, and members of the Allen Institute Neural Coding group for helpful discussions. We thank the Allen Institute founder, Paul G. Allen, for his vision, encouragement and support.

---

**Fig S1.**
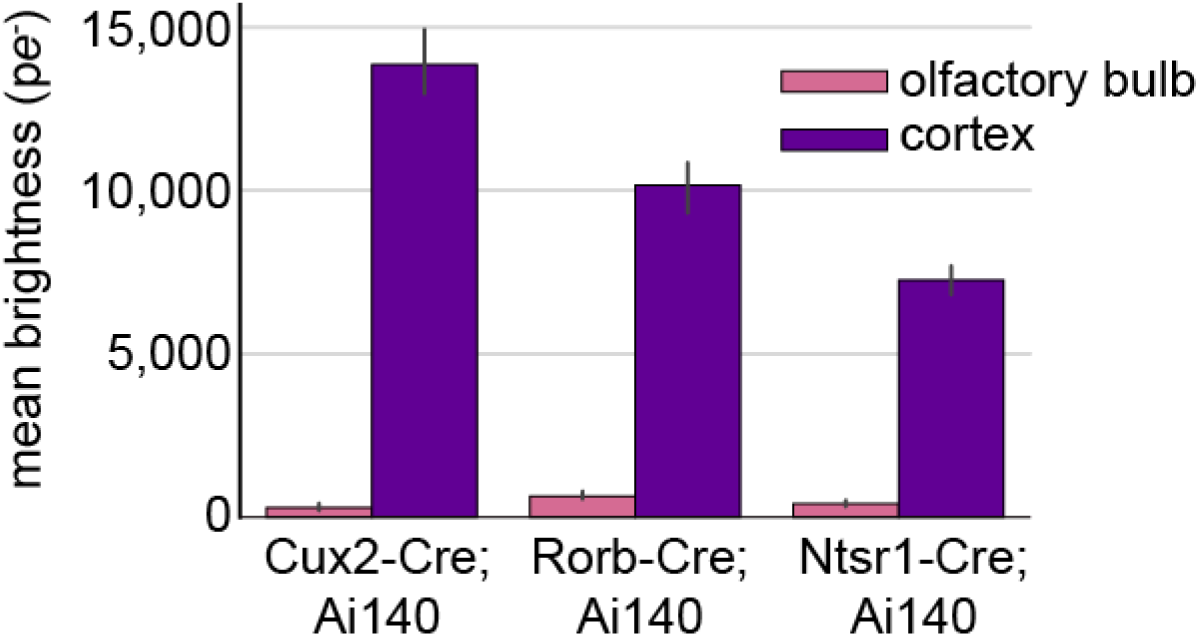
Relative brightness of GFP mice. Image brightness, in mean photoelectrons, calculated from 10×10 pixel regions (65.0 µm^2^) over medial somatosensory cortex from 3 Cux2-Ai140, 3 Rorb-Ai140 and 5 Ntsr1-Ai140 mice. Olfactory bulb measurements were taken to represent the background photon count, including tissue autofluorescence, skull autofluorescence, and light leak.

**Fig S2.**
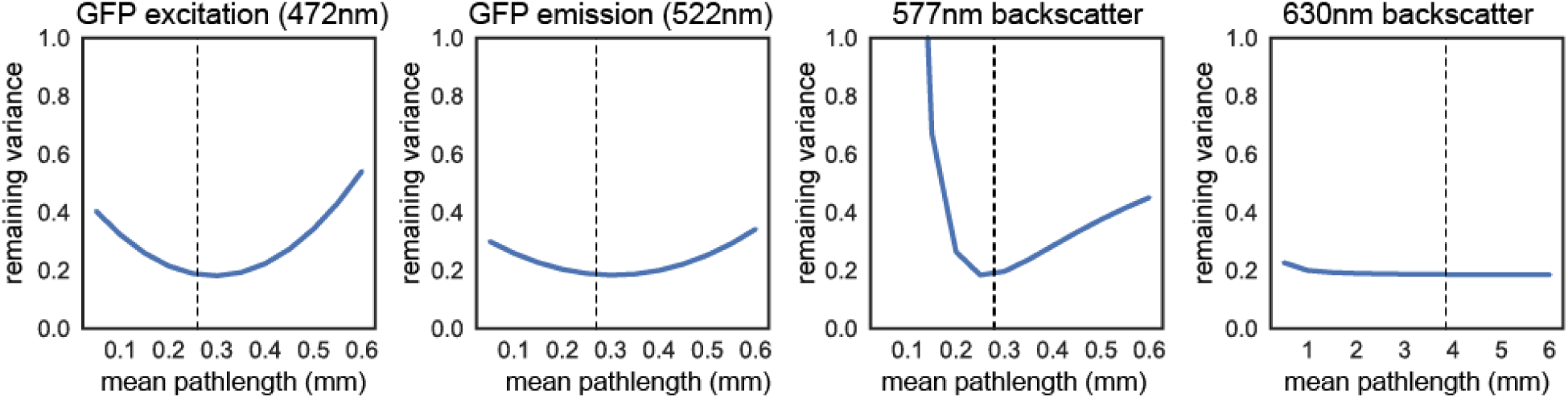
Sensitivity of Beer-Lambert model to changes in path length. The median remaining variance of all pixels across the brain from a single Cux2-Cre;Ai140 mouse while changing one mean path length in the Beer-Lambert model. The sensitivity of model performance to each path length was evaluated with all other parameters (including other path lengths) constant. Path lengths values used elsewhere in the paper are marked with dotted lines: 260µm (GFP excitation), 270µm (GFP emission), 280 µm (577nm backscatter), 3.85mm (630nm backscatter).

**Fig S3.**
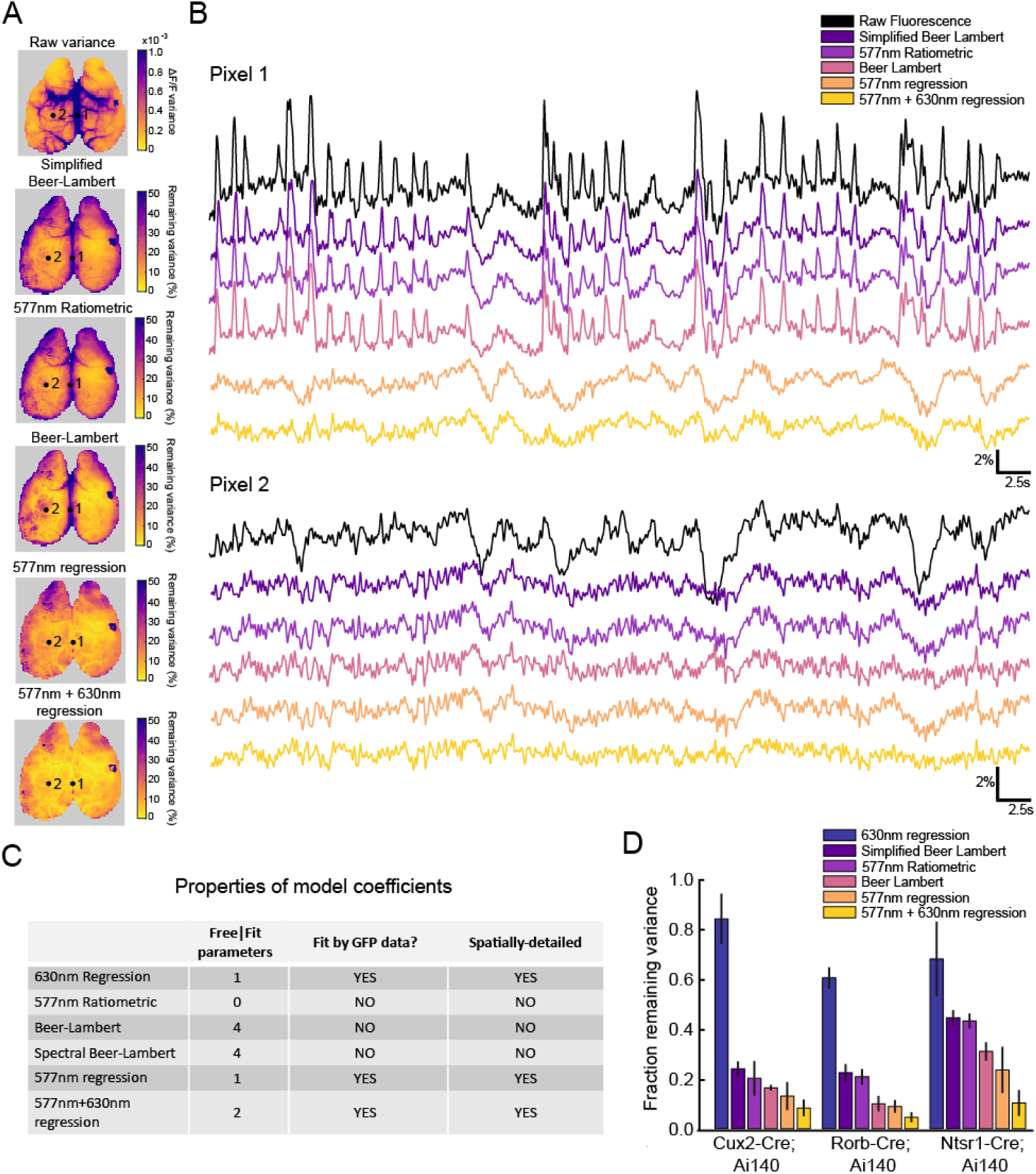
Detailed comparison of all models against GFP results. (A) Map of variance from a Cux2-Ai140 (GFP) mouse (top), and maps of remaining variance after correction using 5 models. (B) Example traces of spontaneous activity and corrected traces from two example pixels (dots, left). Raw data was corrected using five methods: Simplified Beer Lambert (see supplemental appendix), Ratiometric demixing ((1+dF/F) / (1+dR_577nm_/R_577nm_)), Beer-Lambert demixing (see supplemental appendix), 577nm regression (Eq. 9 omitting the term for S_2_), and 577nm + 640nm regression (Eq. 9). (C) Summary of key differences between how coefficients are generated in the five models. (D) Summary of variance remaining after correction (averaged across cortical surface) from three Cre lines. Bars in D represent mean ± STD. 3 Cux2-Ai140 mice, 3 Rorb-Ai140 mice, 5 Ntsr1-Ai140 mice.

**Fig S4.**
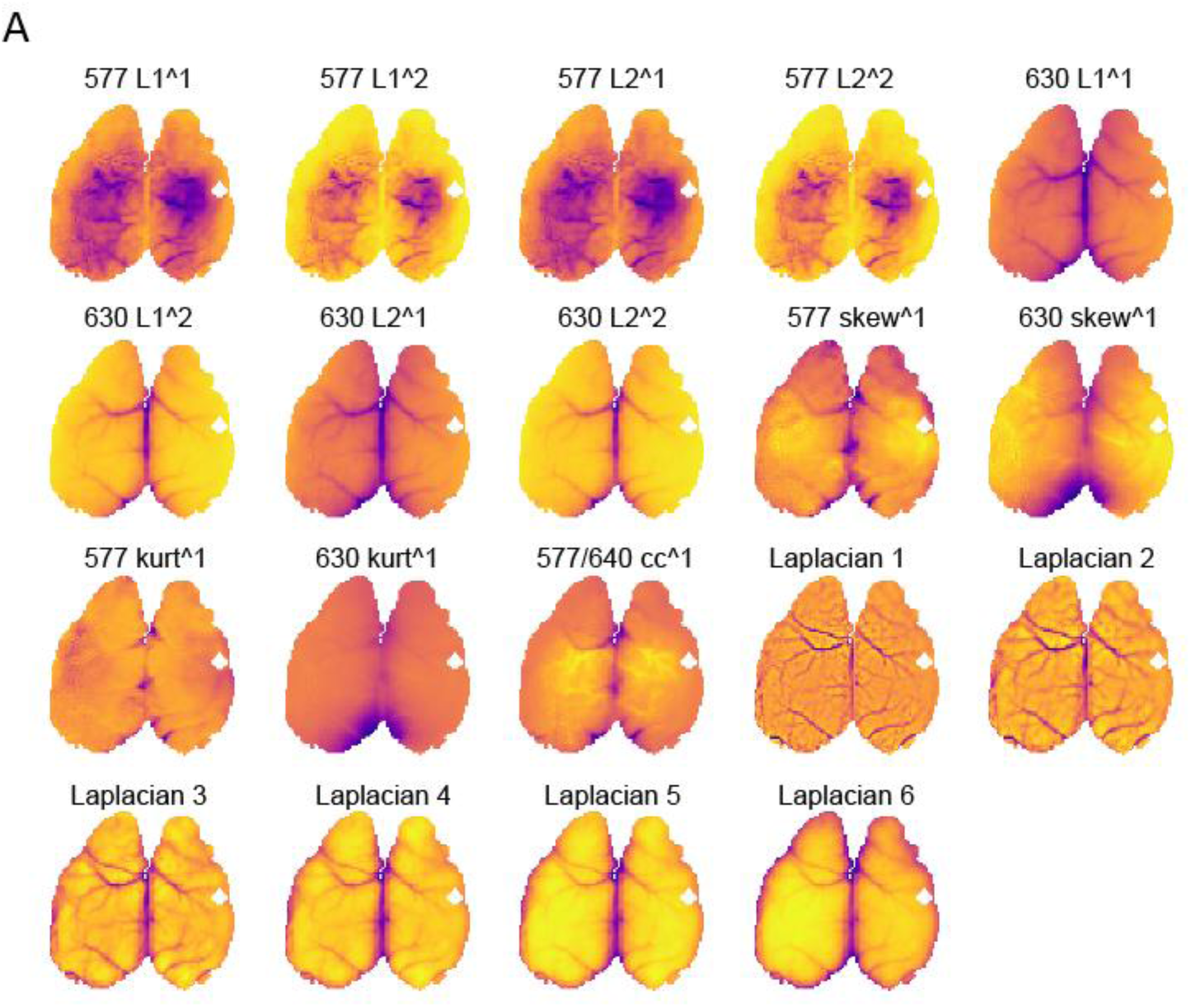
Maps of Spatial Model regressors. (A) 19 different projections of backscatter data (or fluorescence data for Laplacian maps) were used as regressors in the Spatial Model to predict coefficient maps from GFP-expressing mice. Statistical maps are: the L1 norm of ΔI_1_/Ī_1_ and its square, the L2 norm (standard deviation) of ΔI_1_/Ī_1_ and its square, the L1 norm of ΔI_2_/Ī_2_ and its square, the L2 norm of ΔI_2_/Ī_2_ and its square, the skewness of the 577nm and 630nm signals, the kurtosis of the 577nm and 630nm signals, the covariance of ΔI_1_/Ī_1_ and ΔI_2_/Ī_2_, the Laplacian maps. Images from a Cux2-Ai140 mouse.

**Supplementary Table 1.**
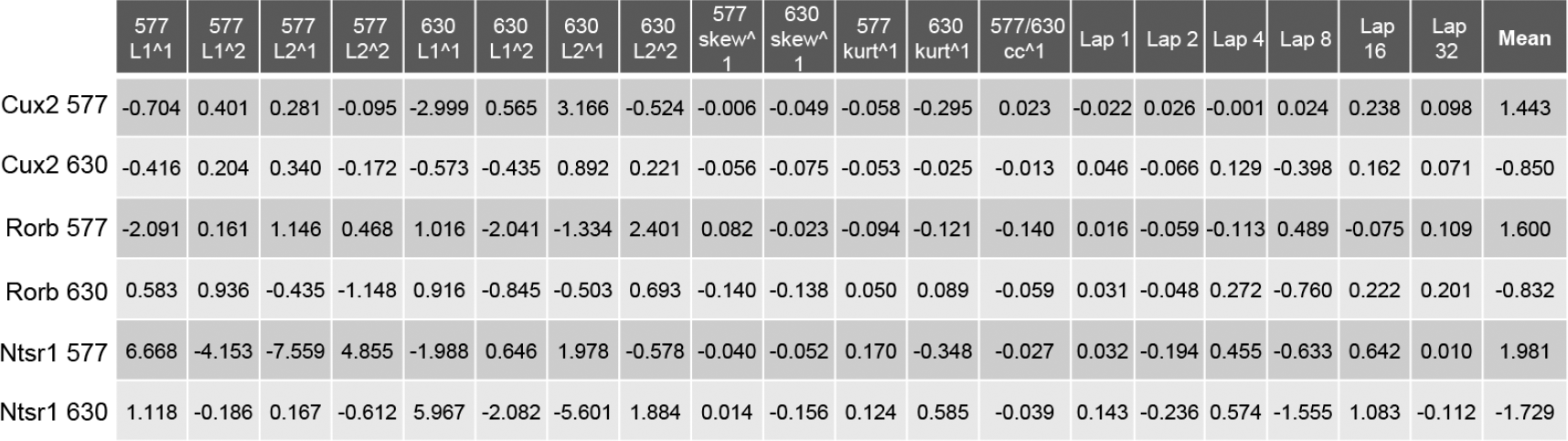
Tabulated coefficients for the Spatial Model. Weights for all spatial-coefficients, each representing the mean weight for the mouse line.

**Fig S5.**
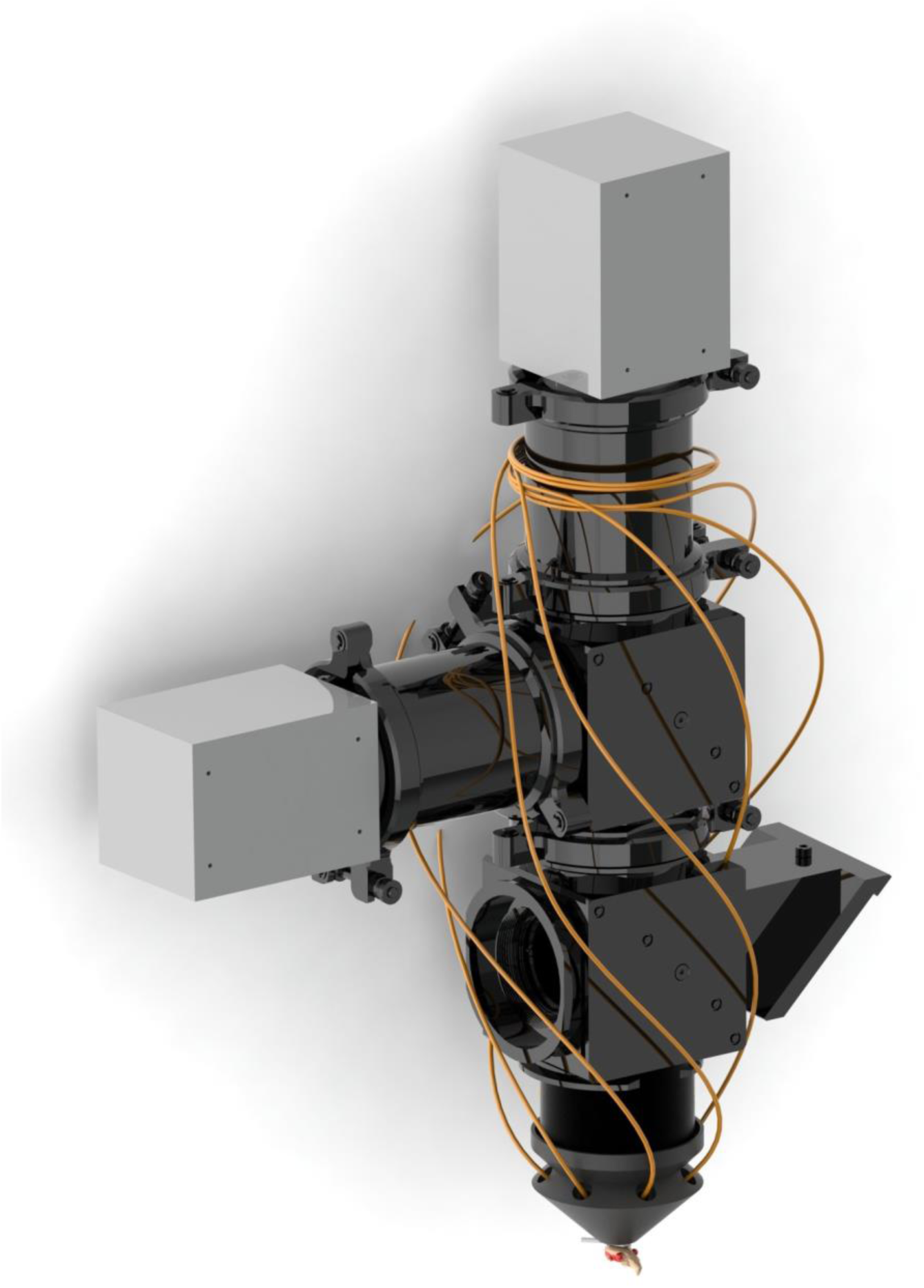
Custom widefield optics system. A modular filter-cube was designed to align cameras, objectives and 2” optics (see also fig 2C for schematic of the optical system). Radial off-axis illumination of mouse cortex for backscatter imaging used a custom light-shield receiving seven LED-coupled optical fibers (LEDs not shown), while epiflourescence excitation enters through the open port in the bottom cube.

### APPENDIX

#### Derivation of the Beer-Lambert and Regression models

A coarse-grained hemodynamic model for the fluorescence intensity recorded from a single pixel, *I*_*F*_(*t*), based on the Beer-Lambert Law and general principles of incoherent optical processes, can be written in compact form as

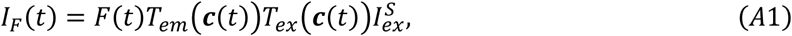

where *F*(*T*) is the coarse-grained intrinsic activation level of the fluorophores averaged over the PSF of the pixel, *T*_*em*_(***c***(*t*)) is the transmittance factor for the light emitted by the fluorophores, *T*_*ex*_(***c***(*t*)) is the transmittance factor for the excitation light, and 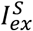 is the excitation source intensity at the specific pixel. The transmittances depend on the oxygenated and deoxygenated hemoglobin concentrations, *C*_*HbO*_(*t*) and *C*_*HbR*_(*t*), respectively, can be combined into a 2-component vector, ***c***(*t*) = [*C*_*HbO*_(*t*), *C*_*HbR*_(*t*)]. Likewise, the recorded intensity of the *b*^*th*^ backscatter channel can be expressed

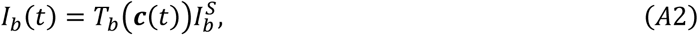

where *T*_*b*_(***c***(*t*)) is the round-trip transmittance, and 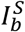 is the corresponding source intensity.

The transmittance factors, *T*_μ_(***c***(*t*)), where μ ∈ {*em, ex, b*}. can be written as

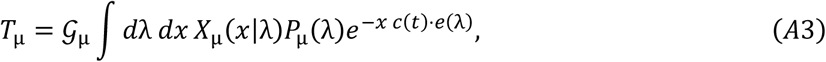

where 𝒢_μ_ contains any geometric factors for the corresponding process, *X*_μ_(*x*|λ) is the distribution over optical path lengths for a given wavelength, λ, and *P*_μ_(λ) is the distribution over wavelengths for the process, including source spectra, filters, and fluorophore excitation/emission spectra, as required. While traveling along a given path, the light is subject to absorption by hemoglobin, in accordance with the Beer-Lambert Law. This is described by the exponential factor in equation 3, which contains the path-length, *x*, the hemoglobin concentrations, *c*(*t*) and *e*(λ) = [*E*_*HbO*_(λ), *E*_*HbR*_(λ)], a two-component vector containing the corresponding hemoglobin molar-extinction coefficients for wavelength λ.

The model has 3 dynamic variables, *F*(*t*), *C*_*HbO*_(*t*), and *C*_*HbR*_(*t*), which can be expressed in terms of mean-values and deviations as 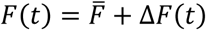 and 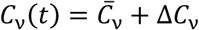, where ν ∈ {*HbO, HbR*}. In the limit where the deviations are small compared to the mean values, as is often the case with wide-field calcium imaging using bright indicator dyes, we can linearize the equations relating the observed intensities to the intrinsic variables. Linearization permits simplification by eliminating some variables. To linearize equations (1) and (2), we first express the intensities as a mean value plus deviation, *I*_*F*_(*t*) = Ī_*F*_ + Δ*I*_*F*_(*t*), and *I*_*b*_(*t*) = Ī_*b*_ + Δ*I*_*b*_(*t*). This leads to

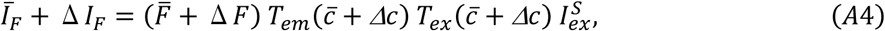

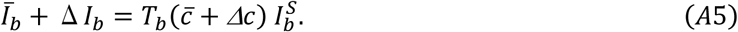

The transmittances can be expanded to first-order in terms of the hemoglobin fluctuations as

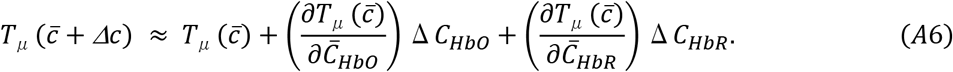

Inserting this expansion into equations (4) and (5), and equating the zeroth order terms with respect to the fluctuations gives the mean intensities

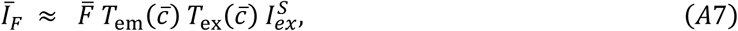

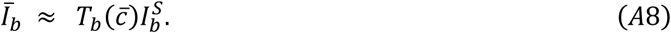

Similarly, equating first-order terms, and dividing by the mean gives

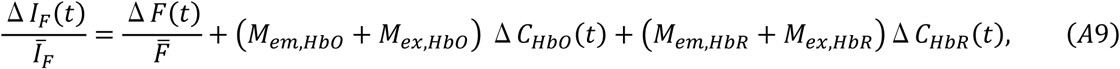

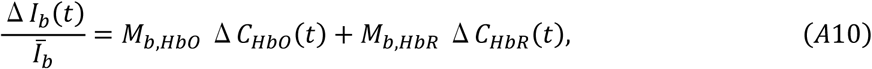

With

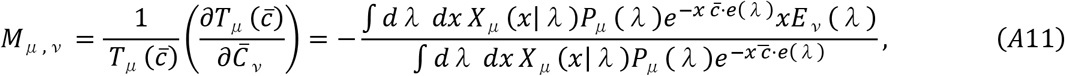

where again *μ* ∈ {*em, ex, b*} and ν ∈ {HbO, HbR}.

The spectral distributions for the light-sources for the fluorescence indicators and for the camera acceptance filters are readily obtained, as are the hemoglobin molar extinction coefficients. The path length distributions and mean hemoglobin molar concentrations, on the other hand, can vary from pixel to pixel and would need to be measured or estimated for each pixel in the field of view.

Because there are two independent hemoglobin components, a minimum of two reflectance channels are necessary to separate the fluorescence from hemodynamics. With a single fluorescence intensity, *I*_*F*_(*t*), and two backscatter channels, *I*_1_(*t*) and *I*_2_(*t*), equation (5) then becomes a pair of backscatter equations,

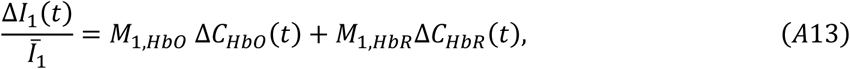

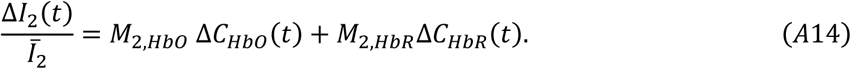

These equations can be solved for the hemodynamic variables, giving

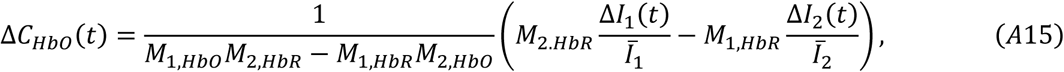

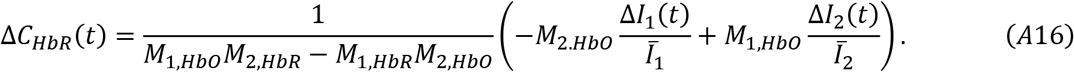

These results can be inserted into the equation for ΔI_*F*_/Ī^*F*^, giving

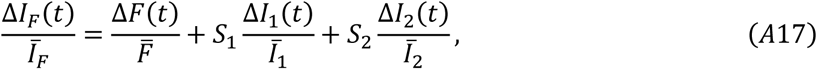

where the coefficients are given by

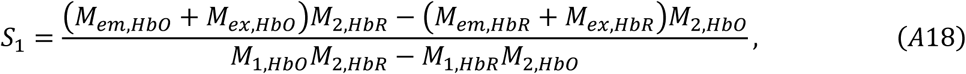

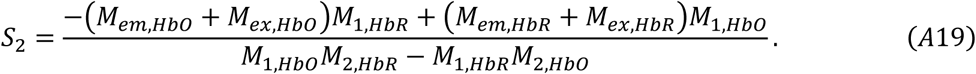

Extending this formalism to the entire image, we arrive at the final formula

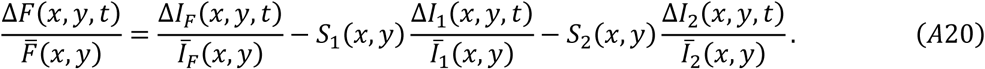

The coefficients, *S*_1_ and *S*_2_, generalize to maps, *S*_1_(*x, y*) and *S*_2_(*x, y*), because the mean hemoglobin concentrations and optical path-length distributions can in principle vary from pixel to pixel.

#### Simplified Beer-Lambert Model: path length and wavelength approximation

There are two major approximations that can be used to simplify the calculations of the backscatter regression coefficients, *S*_1_ and *S*_2_. The first is to replace the distribution over path lengths with a single characteristic path length, x_C_(λ). This is accomplished by taking X(x|λ) → Δ(x - x_C_(λ)), where Δ(x) is the Dirac delta function. The second approximation is to replace the distribution over wavelengths with a single characteristic wavelength, λ_C_, accomplished by taking P(λ) → Δ(λ - λ_C_).

Taking these two approximations together gives

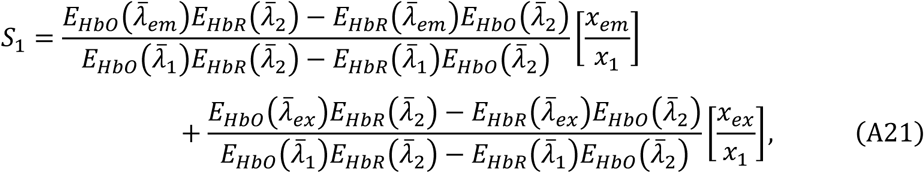

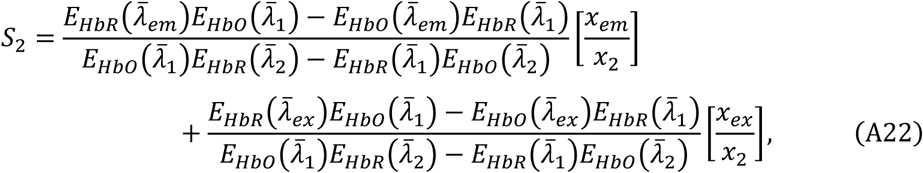

which is the result from (Ma et al. 2016a).

Using our images from GFP mice, we can invalidate this simplification of the Beer-Lambert model (equations 21, 22) for the wavelengths used in our experiment (see figure S3 for remaining variance). Using our wavelengths (λ_ex_ = 473.23 nM, λ_eM_ = 519.99 nM, λ_1_ = 577.20 nM, and λ_2_ = 630.30 nM), and path lengths (577nm: 280 µm, 630nm: 3.85mm, 472nm: 260µm for one-way fluorophore excitation, 522nm: 270µm for one-way fluorescent emission, Ma *et al.*, 2016a), equation (22) predicts a positive value for S_2_ (S_1_ = 1.0314, S_2_ = 0.1289), whereas after regression our data yields a negative value over 99.53% of pixels (see figure 3E). A positive value for S_2_ can only be obtained from Eqs. (21) and (22) for our wavelengths by taking one of the backscatter path lengths to be negative. Hence this simplification of the Beer-Lambert model requires the use of path lengths that are not physically plausible.

#### Spectrally-detailed Beer-Lambert Model

The alternative to the path length and wavelength approximation of the simplified Beer-Lambert model is to replace the path length distributions with single characteristic path lengths (X(x|λ) → Δ(x - x_C_(λ)) equation 11, but to keep the integral over wavelengths and the exact forms of the various spectral distributions *P*_μ_(λ). When applied to our data from GFP mice, with the wavelength integrals computed numerically, this level of approximation results in the correct signs for the backscatter coefficients, *S*_1_ and *S*_2_, so that it is possible to adjust the path-lengths and obtain agreement between the experimental data and the Beer-Lambert model. This is the form of the Beer-Lambert model used in this paper (see figure 3).

To compute the coefficients in this manner requires estimates for the background oxy- and deoxy-hemoglobin concentrations. If the molar concentration of hemoglobin in mouse blood is 2.2 × 10^-3^ mol/L (Raabe *et al.,* 2011), with a typical oxygenation level of 85%, and a Cortical Blood Volume in mouse cortex of ∼0.04 (Chugh *et al.,* 2008), we arrive at *Ć*_*HbO*_ 7.4 × 10^-5^Mol/Land *Ć*_*HbR*_1.3 × 10^-5^Mol/L. We note that the results are insensitive to these numbers, e.g. the dependence vanishes altogether in the approximations (21) and (22). Changing the values over the entire plausible range only changes the resulting coefficients in the third significant figure, hence these reasonable estimates are sufficient.

We apply this to our experimental setup by computing the wavelength integrals in (11) numerically, using directly measured spectra for the two back-scatter channels, combined with manufacturers spectra for our GFP filter, and GFP fluorophore spectra from (http://www.tsienlab.ucsd.edu/Documents.htm). We also use extinction coefficient data from (http://omlc.ogi.edu/spectra/hemoglobin/summary.html), and estimated path-lengths from (Ma *et al.*, 2016a). This leads to the coefficients (S1=1.176, S2=-0.434) which corresponds closely to the regression coefficients calculated in the absence of physical estimates (figure 3E). This shows that a negative 640nm regression coefficient can be obtained from Beer-Lambert with positive path-lengths, and results from the details of the wavelength dependence of the extinction coefficients and path-length data. This is the form of the Beer-Lambert model used in this paper (figure 3).

#### Spatial-Regression Model

Correction of hemodynamics in GCaMP mice presents a challenge for our regression approach because in these animals the fluorescent signal, ΔI_*F*_/ Ī_*F*_, contains an unknown calcium-dependent signal, 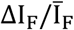 term (Eq. (7), main text). Because of neurovascular coupling, the hemodynamic and calcium-dependent terms are not independent so that a simple regression of ΔI_1_/ Ī_1_ and ΔI_2_/ Ī_2_ from ΔI_*F*_/ Ī_*F*_ will not yield the true calcium dynamic signal.

For a GFP mouse, the coefficient maps, S_1_(x, y) and S_2_(x, y) are obtained by pixel-wise linear regression of the fluorescence signal onto the backscatter signals. To transfer the obtained maps between different mice, we first correlate the features in the maps with corresponding features in the individual brain structures. To do this we constructed a set of *M=19* maps, M_M_(x, y), where M ∈ {1,2, … M}, from the backscatter data and fluorescence data. These maps should be chosen so that they do not reflect calcium dynamics when they are obtained from a GCaMP mouse. Because the data is scaled by the mean-intensities on each channel before any models are constructed, these maps should also be dimensionless and not scale with intensity.

The statistical maps we use are: [1-2] the L1 norm of ΔI_1_/ Ī_1_ and its square, [3-4] the L2 norm of ΔI_1_/ Ī_1_ and its square, [5-6] the L1 norm of ΔI_2_/ Ī_2_ and its square, [7-8] the L2 norm of ΔI_2_/ Ī_2_ and its square, [9-10] the skewness of the 577nm and 630nm signals, [11-12] the kurtosis of the 577nm and 630nm signals, [13] the covariance of ΔI_1_/ Ī_1_ and ΔI_2_/ Ī_2_. To capture the location of the blood-vessels independently of the excitation intensity profile, we compute the ratio of a Gaussian-blurred time-averaged image of the fluorescence emission divided by the time-averaged image itself. Rather than choose a blurring length-scale arbitrarily, we include maps computed at multiple blurring length-scales. The remaining 6 maps are blood-vessel maps computed for length scales of 1, 2, 4, 8, 16, and 32 pixels, respectively (Laplacian maps, figure S4).

We include in our training datasets only those pixels whose fractional variance explained (FVE) is above a chosen threshold of 75%. This is because a low FVE was found to reflect pixels whose initial variance is very low, roughly the same as that obtained on high FVE pixels after demixing, so that the model is fitting noise rather than signal on those pixels.

We constructed three types of Spatial Models: (1) standard Spatial Models, (2) cross-validated Spatial Models, and (3) Cre line Spatial Models.

#### Standard Spatial Model

the 19 maps are z-scored across pixels for a single GFP animal and merged into a 19×D_TRAIN_ predictor array, P_TRAIN_, where D_TRAIN_ is the number of pixels in the training set (i.e. FVE > .75). The two direct-regression coefficient maps, S_1_ and S_2_ for the same set of pixels are merged into a 2×D_TRAIN_ response array, R_TRAIN_, for the same animal. From a linear regression model, R_TRAIN_ - ⟨R_TRAIN_⟩ = C_MM_ - P_TRAIN_, where ⟨…⟩ indicates average over pixels, we obtain the 2×19 coefficient array, C_MM_. We can then reconstruct the response data as R_MM_ = C_MM_P_TEST_+ ⟨R_TRAIN_⟩, where P_TEST_ contains the 19 z- scored predictors from all pixels in the same animal. R_MM_ is then used to demix the fluorescence from the animal, so that the remaining variance can be compared to the optimal result obtained by direct backscatter regression. In this way, we can learn how much error is introduced by the process of transferring the backscatter-regression coefficient maps onto the 19-map representation.

#### Cross-validated Spatial-Model

This model is used to test the amount of error caused by transferring the spatial-model coefficients from one animal to another. For each animal, we again obtain a reconstructed response array, R_MMCV_ = C_MMCV_·P_TEST_+ ⟨R_TRAIN_⟩, but here C_MMCV_ is obtained from a training set consisting of all training pixels (FVE > .75) from all other animals in the same Cre line (i.e. the test animal is excluded from the training data). The testing set then consists of all pixels in the test animal. Here the differential between remaining variance obtained by demixing the fluorescence signal with R- _MMCV_ and that obtained via R_MM_ in a the same GFP mouse estimates the error incurred during transfer of the spatial-model between animals.

#### Cre-line-wise Spatial Model

Here the training set consists of all training pixels (FVE > .75) in all animals in a Cre line. The trained model is then expected to be used to construct the backscatter coefficients for all pixels in a GCaMP mouse. This would represent the best attempt to demix the GCaMP animal using all available training data from the same Cre line. Error limits would come from the cross-validated GFP results, with the caveat that in the case of small cohorts, including the additional animal in the training data will presumably lower the error somewhat, meaning that the cross-validation results are likely an overestimate of the Spatial Model error.

## REFERENCES

Allen, W. E., Kauvar, I. V., Chen, M. Z., Richman, E. B., Yang, S. J., Chan, K., … Deisseroth, K. (2017). Global Representations of Goal-Directed Behavior in Distinct Cell Types of Mouse Neocortex. Neuron, 94(4), 891–907.

Arridge, S. R., Cope, M., & Delpy, D. T. (1992). The theoretical basis for the determination of optical path-lengths in tissue: temporal and frequency analysis. Physics in Medicine and Biology, 37(7), 1531–1560.

Bethge P, Carta S, Lorenzo DA, Egolf L, Goniotaki D, Madisen L, et al.(2017) An R-CaMP1.07 reporter mouse for cell-type-specific expression of a sensitive red fluorescent calcium indicator. PLoS ONE 12(6).

Berwick, J., Johnston, D., Jones, M., Martindale, J., Redgrave, P., McLoughlin, N., … Mayhew, J. E. W. (2005). Neurovascular coupling investigated with two-dimensional optical imaging spectroscopy in rat whisker barrel cortex. European Journal of Neuroscience, 22(7), 1655–1666.

Bouchard, M. B., Chen, B. R., Burgess, S. A., & Hillman, E. M. C. (2009). Ultra-fast multispectral optical imaging of cortical oxygenation, blood flow, and intracellular calcium dynamics. Optics Express, 17(18), 15670.

Chen Q; Cichon J; Wang W; Qiu L; Lee SJ; Campbell NR; DeStefino N; Goard MJ; Fu Z; Yasuda R; Looger LL; Arenkiel BR; Gan WB; Feng G (2012) Imaging neural activity using Thy1-GCaMP transgenic mice. Neuron 76(2), 297–308.

Chugh BP, Lerch JP, Yu LX, Pienkowski M, Harrison RV, Henkelman RM, Sled JG. (2000). Measurement of cerebral blood volume in mouse brain regions using micro-computed tomography. NeuroImage. 47 (4).

Daigle TL, Madisen L et al. (2018). A suite of transgenic driver and reporter mouse lines with enhanced brain cell type targeting and functionality. Cell, July 12.

Dana H, Chen TW, Hu A, Shields BC, Guo C, Looger LL, Kim DS, Svoboda K (2014) Thy1-GCaMP6 transgenic mice for neuronal population imaging in vivo. PLoS One 9.

Devor, A., Sakadžic, S., Srinivasan, V. J., Yaseen, M. A., Nizar, K., Saisan, P. A., … Boas, D. A. (2012). Frontiers in Optical Imaging of Cerebral Blood Flow and Metabolism. Journal of Cerebral Blood Flow & Metabolism, 32(7).

Franco SJ, Gil-Sanz C, Martinez-Garay I, Espinosa A, Harkins-Perry SR, Ramos C, Müller U (2012) Fate-restricted neural progenitors in the mammalian cerebral cortex. Science 337(6095), 746–749.

Frostig, R. D., Lieke, E. E., Ts’o, D. Y., & Grinvald, A. (1990). Cortical functional architecture and local coupling between neuronal activity and the microcirculation revealed by in vivo high-resolution optical imaging of intrinsic signals. Proceedings of the National Academy of Sciences, 87(16), 6082–6086.

Gilad, A., Gallero-Salas, Y., Groos, D., & Helmchen, F. (2018). Behavioral Strategy Determines Frontal or Posterior Location of Short-Term Memory in Neocortex. Neuron, 99(4), 814–828.

Gisolf, J., Van Lieshout, J. J., Van Heusden, K., Pott, F., Stok, W. J., & Karemaker, J. M. (2004). Human cerebral venous outflow pathway depends on posture and central venous pressure. The Journal of Physiology, 560(1), 317–327.

Gong S, Doughty M, Harbaugh CR, Cummins A, Hatten ME, Heintz N, Gerfen CR (2007) Targeting Cre recombinase to specific neuron populations with bacterial artificial chromosome constructs. J Neurosci 27(37), 9817–9823.

Herman, M. C., Cardoso, M. M. B., Lima, B., Sirotin, Y. B., & Das, A. (2017). Simultaneously estimating the task-related and stimulus-evoked components of hemodynamic imaging measurements. Neurophotonics, 4(3).

Hillman, E. M. C., Devor, A., Bouchard, M. B., Dunn, A. K., Krauss, G. W., Skoch, J., … Boas, D. A. (2007). Depth-resolved optical imaging and microscopy of vascular compartment dynamics during somatosensory stimulation. NeuroImage, 35(1).

Hillman, E. M. C. (2014). Coupling Mechanism and Significance of the BOLD Signal: A Status Report. Annual Review of Neuroscience, 37(1), 161–181.

Harris, J. A., Hirokawa, K. E., Sorensen, S. A., Gu, H., Mills, M., Ng, L. L., … Zeng, H. (2014). Anatomical characterization of Cre driver mice for neural circuit mapping and manipulation. Frontiers in Neural Circuits.

Hasan MT, Friedrich RW, Euler T, Larkum ME, Giese G, Both M, Duebel J, Waters J, Bujard H, Griesbeck O, Tsien RY, Nagai T, Miyawaki A, Denk W (2004) Functional fluorescent Ca2+ indicator proteins in transgenic mice under TET control. PLoS Biol 2, 763–75.

Huo, B.-X., Gao, Y.-R., & Drew, P. J. (2015). Quantitative separation of arterial and venous cerebral blood volume increases during voluntary locomotion. NeuroImage, 105, 369–379.

Kalatsky, V. A., & Stryker, M. P. (2003). New Paradigm for Optical Imaging. Neuron, 38(4), 529–545.

Kohl, M., Lindauer, U., Royl, G., Kühl, M., Gold, L., Villringer, A., & Dirnagl, U. (2000). Physical model for the spectroscopic analysis of cortical intrinsic optical signals. Physics in Medicine and Biology, 45(12), 3749–3764.

Logothetis, N. K., & Wandell, B. A. (2004). Interpreting the BOLD Signal. Annual Review of Physiology, 66(1), 735–769.

Ma, Y., Shaik, M. A., Kim, S. H., Kozberg, M. G., Thibodeaux, D. N., Zhao, H. T., … Hillman, E. M. C. (2016). Wide-field optical mapping of neural activity and brain haemodynamics: considerations and novel approaches. Philosophical Transactions of the Royal Society B: Biological Sciences, 371(1705).

Ma, Y., Shaik, M. A., Kozberg, M. G., Kim, S. H., Portes, J. P., Timerman, D., & Hillman, E. M. C. (2016). Resting-state hemodynamics are spatiotemporally coupled to synchronized and symmetric neural activity in excitatory neurons. Proceedings of the National Academy of Sciences, 113(52).

Madisen L, Garner AR, Shimaoka D, Chuong AS, Klapoetke NC, Li L, van der Bourg A, Niino Y, Egolf L, Monetti C, Gu H, Mills M, Cheng A, Tasic B, Nguyen TN, Sunkin SM, Benucci A, Nagy A, Miyawaki A, Helmchen F, et al.(2015) Transgenic mice for intersectional targeting of neural sensors and effectors with high specificity and performance. Neuron 85, 942–958.

Malonek, D., & Grinvald, A. (1996). Interactions Between Electrical Activity and Cortical Microcirculation Revealed by Imaging Spectroscopy: Implications for Functional Brain Mapping. Science, 272(5261), 551–554.

Makino H, Ren C, Liu H, Kim AN, Kondapaneni N, Liu X, Kuzum D, Komiyama T (2017) Transformation of cortex-wide emergent properties during motor learning. Neuron 94(4), 880–890.

Mitra, A., Kraft, A., Wright, P., Acland, B., Snyder, A. Z., Rosenthal, Z., … Raichle, M. E. (2018). Spontaneous Infra-slow Brain Activity Has Unique Spatiotemporal Dynamics and Laminar Structure. Neuron, 98(2)

Mohajerani, M. H., Chan, A. W., Mohsenvand, M., LeDue, J., Liu, R., McVea, D. A., … Murphy, T. H. (2013). Spontaneous cortical activity alternates between motifs defined by regional axonal projections. Nature Neuroscience, 16(10).

O’Herron, P., Chhatbar, P. Y., Levy, M., Shen, Z., Schramm, A. E., Lu, Z., & Kara, P. (2016). Neural correlates of single-vessel haemodynamic responses in vivo. Nature, 534(7607), 378–382.

Qvarlander, S., Sundström, N., Malm, J., & Eklund, A. (2013). Postural effects on intracranial pressure: modeling and clinical evaluation. Journal of Applied Physiology, 115(10), 1474–1480.

Raabe BM, Artwohl JE, Purcell JE, Lovaglio J, Fortman JD. (2011) Effects of Weekly Blood Collection in C57BL/6 Mice. J Am. Assoc. Lab. Anim. Sci. 50(5).

Silasi G, Xiao D, Vanni MP, Chen AC, Murphy TH (2016) Intact skull chronic windows for mesoscopic wide-field imaging in awake mice. J Neurosci Methods 267, 141–149.

Sirotin, Y. B., Hillman, E. M. C., Bordier, C., & Das, A. (2009). Spatiotemporal precision and hemodynamic mechanism of optical point spreads in alert primates. Proceedings of the National Academy of Sciences, 106(43).

Steinmetz NA, Buetfering C, Lecoq J, Lee CR, Peters AJ, Jacobs EAK, Coen P, Ollerenshaw DR, Valley MT, de Vries SEJ, Garrett M, Zhuang J, Groblewski PA, Manavi S, Miles J, White C, Lee E, Griffin F, Larkin JD, Roll K, Cross S, Nguyen TV, Larsen R, Pendergraft J, Daigle T, Tasic B, Thompson CL, Waters J, Olsen S, Margolis DJ, Zeng H, Hausser M, Carandini M, Harris KD (2017) Aberrant cortical activity in multiple GCaMP6-expressing transgenic mouse lines. eNeuro 4(5).

Takatani S, Graham MD. (1979). Theoretical analysis of diffuse reflectance from a two-layer tissue model. IEEE Trans Biomed Eng. Dec 26 (2).

Wekselblatt JB; Flister ED; Piscopo DM; Niell CM (2016) Large-scale imaging of cortical dynamics during sensory perception and behavior. J Neurophysiol 115(6), 2852–2866.

White, B. R., Bauer, A. Q., Snyder, A. Z., Schlaggar, B. L., Lee, J.-M., & Culver, J. P. (2011). Imaging of Functional Connectivity in the Mouse Brain. PLoS ONE, 6(1)

Winder, A. T., Echagarruga, C., Zhang, Q., & Drew, P. J. (2017). Weak correlations between hemodynamic signals and ongoing neural activity during the resting state. Nature Neuroscience, 20(12), 1761–1769.

Wray S, Cope M, Delpy DT, Wyatt JS, Reynolds EO. (1988). Characterization of the near infrared absorption spectra of cytochrome aa3 and haemoglobin for the non-invasive monitoring of cerebral oxygenation. Biochim Biophys Acta. March 30.

Xiao, D., Vanni, M. P., Mitelut, C. C., Chan, A. W., LeDue, J. M., Xie, Y., … Murphy, T. H. (2017). Mapping cortical mesoscopic networks of single spiking cortical or sub-cortical neurons. eLife.

Zariwala HA, Borghuis BG, Hoogland TM, Madisen L, Tian L, De Zeeuw CI, Zeng H, Looger LL, Svoboda K, Chen TW (2012) A Cre-dependent GCaMP3 reporter mouse for neuronal imaging in vivo. J Neurosci 32(9), 3131–3141.

Zipfel, W. R., Williams, R. M., Christie, R., Nikitin, A. Y., Hyman, B. T., & Webb, W. W. (2003). Live tissue intrinsic emission microscopy using multiphoton-excited native fluorescence and second harmonic generation. PNAS, 100(12), 7075–7080.

Zhuang J, Ng L, Williams D, Valley M, Li Y, Garrett M, Waters J (2017) An extended retinotopic map of mouse cortex. eLife 6, e18372. PMID28059700.

